# Stabilization versus flexibility: detergent-dependent trade-offs in neurotensin receptor 1 GPCR ensembles

**DOI:** 10.1101/2025.09.22.677830

**Authors:** James B. Bower, Wijnand J. C. van der Velden, Karen P Gomez, Mingzhe Pan, Fabian Bumbak, Nagarajan Vaidehi, Joshua J. Ziarek

## Abstract

Detergents provide essential membrane-mimetic environments for studying G protein-coupled receptors (GPCRs), but their molecular impact on receptor energetics remains incompletely understood. We combined ligand binding, thermostability measurements and atomistic molecular dynamics to dissect detergent- versus ligand-driven stabilization in a thermostabilized neurotensin receptor 1 (enNTS1). Circular dichroism and ligand binding assays revealed that apo enNTS1 becomes progressively more stable in decyl maltoside (DM), dodecyl maltoside (DDM), and lauryl maltose neopentyl glycol (LMNG). Yet this gain in baseline stability was accompanied by a paradox: LMNG, the most stabilizing detergent, supported the weakest neurotensin agonist binding affinity. Thermodynamic analysis resolved this contradiction by partitioning stability into detergent-driven conformational rigidity (ΔG_conf_) and ligand-induced stabilization (ΔG_ligand_). In DM, ΔG_ligand_ contributions were large, consistent with the receptor’s engineered background. In contrast, LMNG maximized ΔG_conf_, constraining conformational flexibility and reducing ΔG_ligand_. Molecular dynamics simulations corroborated these results, showing that LMNG formed denser, less mobile detergent shells around the receptor, enhancing protein–detergent interaction energies while limiting conformational flexibility. Redistribution of ligand contacts, particularly at neurotensin residue Y11, further underscored detergent-dependent modulation of the binding pocket. These results highlight a fundamental trade-off: LMNG provides exceptional receptor stabilization, supporting structural studies, but may mask conformational states relevant to signaling. In contrast, less rigid detergents preserve ligand-induced transitions at the expense of stability. These findings emphasize that detergent choice should be guided by whether the goal is structural resolution or dynamic characterization.

## INTRODUCTION

G protein-coupled receptors (GPCRs) constitute the largest family of plasma membrane receptors in eukaryotes, functioning as versatile sensors of hormones, neurotransmitters, peptides, and metabolites^1,2^. Their pharmacological importance is underscored by the fact that ∼34% of approved drugs target GPCRs, spanning indications from cardiovascular to neurological disorders^3,4^. Despite this therapeutic relevance, high-resolution structures exist for fewer than 200 unique receptors out of the more than 800 encoded in the human genome^5,6^. A major challenge remains the purification and stabilization of GPCRs in membrane-mimetic environments that maintain native conformational equilibria^7^.

Although nanodiscs, bicelles, and liposomes have expanded the membrane-mimetic toolkit^8,9^, detergent micelles remain indispensable for initial solubilization and purification. Nonionic maltoside detergents such as n-decyl-β-D-maltoside (DM) and n-dodecyl-β-D-maltoside (DDM) have been widely used owing to their relatively mild properties^10^. More recently, branched maltose-neopentylglycol (MNG) detergents were introduced to enhance receptor stability and crystallization success^11^. Among these, lauryl maltose neopentyl glycol (LMNG) has emerged as a workhorse, used to solve more than half of all GPCR structures^12^. LMNG improves thermostability in multiple receptors, including adenosine A2A receptor (A2AR), β2-adrenergic receptor (β2AR), opioid receptors, and muscarinic receptors^13-23^. LMNG’s branched design yields tighter micelle packing, reduced detergent mobility, and greater thermal stabilization^24,25^. However, the energetic consequences of this stabilization are incompletely understood, particularly regarding how detergents redistribute the balance between conformational rigidity and ligand-induced effects.

Ligand binding is known to enhance GPCR stability by increasing global rigidity^26^, yet this contribution (ΔG_ligand_) depends on the baseline conformational ensemble established by the membrane-mimetic environment. A detergent that rigidifies the apo state may maximize conformational stability (ΔG_conf_) but simultaneously reduce the conformational freedom available for ligand-induced stabilization. Such trade-offs have direct implications for receptor activation, as conformational shifts between apo and ligand-bound states underlie GPCR signaling^27,28^.

Here, we address this question by dissecting detergent versus ligand contributions in enNTS1, a thermostabilized variant of the rat neurotensin receptor 1. enNTS1 was originally selected for stability in DM detergent^29^, providing an ideal background for testing how DM, DDM, and LMNG redistribute stabilization. Using CD spectroscopy, ligand-binding thermostability assays, and atomistic molecular dynamics, we show that LMNG maximizes ΔG_conf_ at the expense of ΔG_ligand_. The paradoxical result—highest baseline stability but weakest agonist affinity—is explained by LMNG’s ability to restrict conformational heterogeneity and remodel ligand contact profiles (notably at NT(8-13) residue Y11)^27,28,30^. This partitioning framework provides a generalizable lens for understanding how membrane mimetics alter GPCR energetics, ligand recognition, and activation pathways^31-33^.

## RESULTS

### Detergent environments differentially influence agonist binding affinity of enNTS1

Detergents are well known to influence GPCR stability, yet their effect on ligand binding affinity has received far less attention. To directly compare micelle environments, we measured the affinity of NT(8-13), the minimal activating peptide of neurotensin, for enNTS1 solubilized in DM, DDM, or LMNG micelles using biolayer interferometry (BLI). NT(8-13) was immobilized via an N-terminal hexaHis tag on the BLI probe, and binding was measured against increasing receptor concentrations. As illustrated in Figure 1, steady-state analysis yielded K_d_ values of 13.7 ± 3.3 nM (DM), 35.9 ± 5.1 nM (DDM), and 94.9 ± 26 nM (LMNG). The highest affinity in DM is expected, given that the enNTS1 construct was thermostabilized for improved NT(8-13) binding in this detergent environment^29^. More surprising, however, is that LMNG, which is predicted to provide greater overall receptor stability than DDM, supports weaker binding affinity. This paradox suggests that detergent environments influence not only receptor thermostability but also the relative energetic contribution of ligand binding. To explore this possibility, we next examined the thermostability of apo and ligand-bound enNTS1 across the same detergent series.

**Figure 1.**
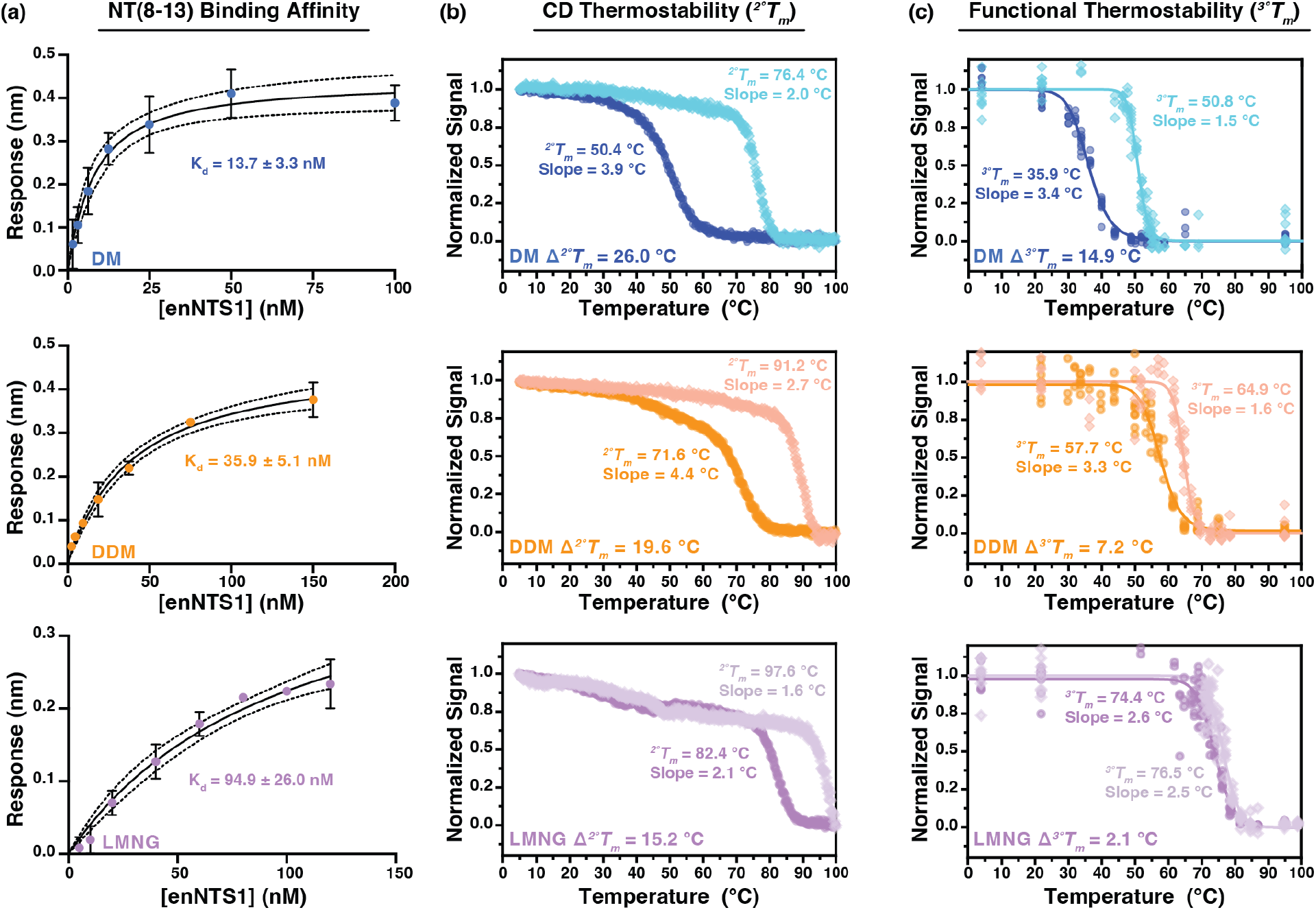
Detergent-dependent affinity and thermostability of enNTS1. (a) NT(8-13) was immobilized through an N-terminal hexaHis tag on the BLI probe, and binding was measured against increasing receptor concentrations solubilized in DM (top), DDM (middle), or LMNG (bottom) micelles. Equilibrium dissociation constants (K_d_) were derived by fitting steady-state kinetics to a quadratic binding model. 95% confidence bands are illustrated as dashed lines above/below the fitted curve. (b) Far-UV CD spectra were collected at increased temperatures for 5 µM enNTS1 in DM (top), DDM (middle), and LMNG (bottom). Apo and NT(8-13)-bound enNTS1 are colored darker and lighter, respectively. The absorption at 222 nm was normalized and fit with a six-parameter Boltzmann equation to yield apparent secondary structure melting temperature (^*2°*^*T*_*m*_) and slope ^36^. (c) Apparent tertiary structure melting temperature (^*3°*^*T*_*m*_) curves monitor the ability of enNTS1 to bind Alexa Fluor 647 dye-labeled NT(8-13) at increasing temperatures. Measurements were collected with 5 µM enNTS1 in DM (top), DDM (middle), and LMNG (bottom) in both apo (darker color) and NT(8-13)-bound (lighter color) states. The data was normalized and fit with a six-parameter Boltzmann equation to yield ^*3°*^*T*_*m*_ and slope.

### Detergent environments differentially stabilize apo and ligand-bound enNTS1

Circular dichroism (CD) spectroscopy was used to assess how detergents alter the thermostability of enNTS1 secondary structure. Far-UV CD reports on α-helical content through minima at ∼210 and 222 nm. Monitoring ellipticity at 222 nm as a function of temperature provided melting curves from which we determined the apparent secondary structure melting temperature (^*2°*^*T*_*m*_). Measurements were performed in DM, DDM, and LMNG micelles for both apo enNTS1 and the agonist-bound state with NT(8-13). Apo enNTS1 displayed the lowest stability in DM micelles (^*2°*^*T*_*m*_ = 50.4 ± 0.07 °C), which increased to 71.6 ± 0.2 °C in DDM and 82.4 ± 0.1 °C in LMNG (Figure 1b). NT(8-13) binding increased the ^*2°*^*T*_*m*_ in all detergents, yielding values of 76.4 ± 0.06 °C, 91.2 ± 0.2 °C, and 97.6 ± 0.2 °C for DM, DDM, and LMNG, respectively (Figure 1b). Ligand-induced stability is not unusual within the GPCR superfamily^26^ and it was shown previously that NT(8-13) binding increases the global rigidity of DDM reconstituted enNTS1^34^. Because enNTS1 was mutationally thermostabilized for high-affinity NT(8-13) binding in DM, the large change in thermostability between apo and holo conditions (Δ^*2°*^*T*_*m*_) observed in DM (26.0 °C) is expected. More revealing is the trend across detergents: Δ^*2°*^*T*_*m*_ decreased to 19.6 °C in DDM and 15.2 °C in LMNG, despite LMNG conferring the highest overall stability. Melting slopes further emphasized this pattern. In DM, the slope steepened substantially upon ligand binding (apo = 3.9 °C, holo = 2.0 °C), consistent with enhanced unfolding cooperativity. DDM showed a similar but less pronounced effect (apo = 4.4 °C, holo = 2.7 °C). In contrast, LMNG exhibited steep slopes that changed little with ligand addition (apo = 2.1 °C, holo = 1.6 °C), suggesting that detergent-driven rigidity minimized ligand-induced contributions.

The apparent tertiary structure melting temperature (^*3°*^*T*_*m*_) was evaluated by measuring loss of NT(8-13) binding as a function of temperature using Alexa Fluor 647-labeled peptide (Figure 1c). Apo enNTS1 was again least stable in DM micelles (^*3°*^*T*_*m*_ = 35.9 ± 0.2 °C), increasing to 57.7 ± 0.4 °C in DDM and 74.4 ± 0.4 °C in LMNG. NT(8-13) binding increased ^*3°*^*T*_*m*_ in all detergents, with values of 50.8 ± 0.2 °C (DM), 64.9 ± 0.2 °C (DDM), and 76.5 ± 0.3 °C (LMNG). As expected, the enNTS1 tertiary structure is less stable than the individual helices^35^; yet, the relative increases in ligand-mediated stabilization mirrored the secondary structure results: Δ^*3°*^*T*_*m*_ = 14.9 °C in DM, 7.2 °C in DDM, and only 2.1 °C in LMNG. Analysis of tertiary unfolding slopes reinforced these findings. In DM, the slope decreased from 3.4 °C (apo) to 1.5 °C (holo), and in DDM from 3.3 °C (apo) to 1.6 °C (holo), indicating that ligand binding sharpened unfolding cooperativity in both environments. In LMNG, however, slopes were nearly unchanged (apo = 2.6 °C, holo = 2.5 °C), again pointing to detergent-imposed rigidity.

Plotting secondary versus tertiary structure melting temperatures revealed a strong correlation for both apo (Slope = 1.18, R^2^ = 0.99) and holo (Slope = 1.17, R^2^ = 0.97) states (Figure S1), consistent with a shared structural basis for unfolding across detergents. Taken together, these results indicate that the large ligand-induced stabilization in DM reflects the engineered background of enNTS1, while the progressively smaller *T*_*m*_ and slope changes in DDM and LMNG emphasize how detergent environments shift the balance of stability contributions. In LMNG, where detergent interactions (ΔG_conf_) dominate, ligand contributions (ΔG_ligand_) are minimized.

### Molecular dynamics models of enNTS1 in detergent micelles

To investigate the mechanisms of enNTS1 stability in different detergents, we performed all atom molecular dynamics (MD) simulations of the receptor in DM, DDM, and LMNG detergents (Table 1). Initial models of the apo and NT(8-13)-bound states were built upon crystal structures of a similar, thermostabilized rat NTS1 (PDB:6Z66 and PDB:6YVR, respectively)^37^. Intracellular loop 3 (ICL3), which was truncated in the crystal constructs, was reintroduced, while the C-terminal DARPin fusion was removed. Helix 8 was modeled from another thermostabilized rat NTS1 structure (PDB: 4BWB)^38^. Finally, the primary sequence of these homology models was modified to match the receptor construct used in the thermostability experiments. Receptor–detergent complexes were generated using the CHARMM-GUI Micelle Builder^39,40^. Each of the six systems (apo and NT(8-13)-bound states in DM, DDM, and LMNG) was simulated in five independent 1 μs trajectories with randomized initial velocities.

**Table 1.** MD parameters for enNTS1 in different detergents. Starting conditions for the CHARMM-based enNTS1 models in DM, DDM, and LMNG detergent micelles.

| **Simulation** | **Base Structure (PDB)** | **Helix 8 Modeling (PDB)** | **Detergent Molecules** | **Water Molecules** | **NaCl (mM)** |
| --- | --- | --- | --- | --- | --- |
| DM Apo | 6Z66 | 4BWB | 192 | 35,445 | 150 |
| DDM Apo | 6Z66 | 4BWB | 192 | 35,206 | 150 |
| LMNG Apo | 6Z66 | 4BWB | 88 | 49,957 | 150 |
| DM NT(8-13) | 6YVR | 4BWB | 192 | 39,586 | 150 |
| DDM NT(8-13) | 6YVR | 4BWB | 192 | 39,364 | 150 |
| LMNG NT(8-13) | 6YVR | 4BWB | 88 | 54,745 | 150 |

Structural stability was assessed by RMSD (root-mean-square deviation) from the starting structure (Figure S2). Percent helicity was then plotted as a function of TM backbone RMSD from the initial structure (Figure S3). Apo models maintained ∼2.5–5% higher helicity than their starting crystal structures, with fluctuations most pronounced in TM5. In the apo state, LMNG-solubilized enNTS1 showed a slightly lower average TM backbone RMSD compared to DM and DDM, but with a broader distribution, suggesting greater conformational heterogeneity. NT(8-13) binding reduced helicity, consistent with crystal structures, and decreased average TM backbone RMSD across all detergent conditions. Backbone root-mean-square fluctuations (RMSF) mapped onto representative structures revealed similar dynamic regions across detergents, with apo receptors particularly flexible in the extracellular region of TM7 and the intracellular portions of TM5/6 (Figure S3). Ligand binding suppressed extracellular fluctuations around the orthosteric pocket, indicating that agonist engagement quenches local flexibility regardless of detergent environment.

### Molecular dynamics reveal detergent-dependent energetic partitioning in enNTS1

To assess how detergent environments shape receptor energetics, we calculated total receptor–detergent interaction energies at 50, 200, 400, and 1000 ns for each MD trajectory. These values included Lennard-Jones and Coulomb nonbonded energies between the enNTS1 transmembrane backbone and surrounding detergent molecules (Figure S4). In the apo state, total interaction energies increased progressively from DM to DDM to LMNG. At 1000 ns, the average interaction energy rose by ∼30 kJ/mol from DM to DDM and by an additional ∼38 kJ/mol from DDM to LMNG (Figure S4a). This trend parallels the experimental thermostability data, consistent with LMNG forming the most stabilizing detergent–protein interactions. In the NT(8-13)-bound state, DM again had the lowest interaction energy, but DDM exceeded LMNG (Figure S4b). Relative to apo, the increase in interaction energy was ∼48 kJ/mol in DM, ∼83 kJ/mol in DDM, and only ∼28 kJ/mol in LMNG. Thus, the apo-to-holo energetic shift was largest in DDM and smallest in LMNG.

This outcome mirrors the thermostability measurements: Δ*T*_*m*_ values were greatest in DM, intermediate in DDM, and minimal in LMNG. Importantly, the large Δ*T*_*m*_ in DM is expected given that the enNTS1 construct was thermostabilized based on NT(8-13) binding in DM. Plotting the calculated total internal energies against the experimentally determined thermostability values produces linear correlations with *R*^*2*^ = 0.88 and 0.65 for secondary and tertiary structure thermostabilities, respectively (Figure 2). Interestingly, removing the outlying DDM NT(8-13) condition increases the correlation to an *R*^*2*^ value of 0.97 for the CD and 0.85 for the functional thermostability data; this datapoint was not removed as there is no justification for its elimination. The paradox therefore emerges most clearly when comparing LMNG and DDM: LMNG provides greater baseline stability, yet ligand-induced energetic stabilization is smaller. In terms of the thermodynamic framework, LMNG favors detergent-driven stabilization (ΔG_conf_), while DDM permits a larger contribution from ligand binding (ΔG_ligand_).

**Figure 2.**
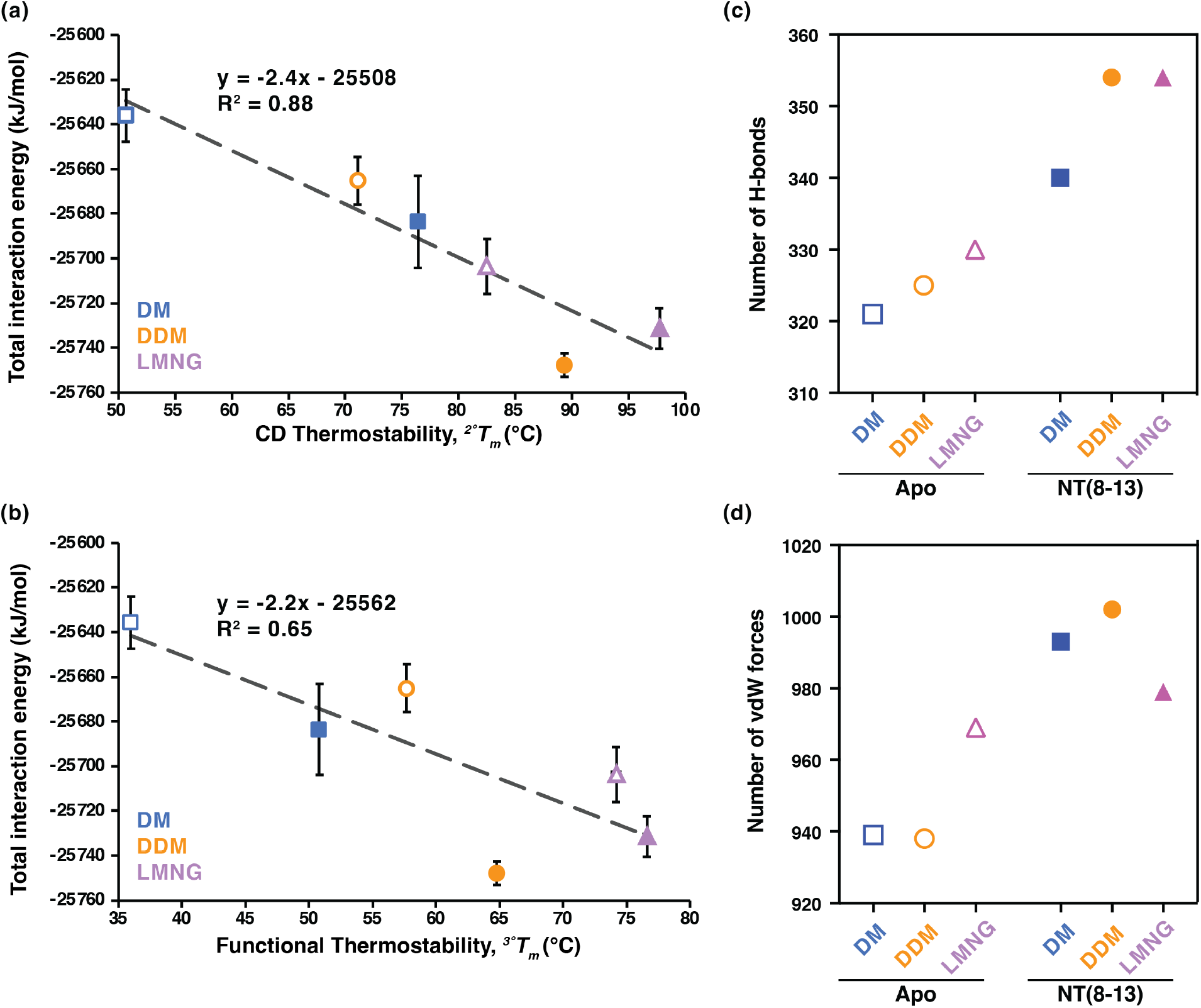
Detergent-dependent interaction energies reveal energetic partitioning of enNTS1 stability. Plots of the MD total interaction energy against (a) circular dichroism and (b) functional thermostability measurements yields linear correlations. enNTS1 in DM (blue), DDM (orange), and LMNG (purple) detergent conditions are plotted for both the apo (open shape) and NT(8-13)-bound (filled shape) states. The number of interhelical (c) hydrogen bonds and (d) van der Waals forces for apo and NT(8-13)-bound enNTS1 in DM, DDM, and LMNG. Analysis was performed on the 5 μs (N= 5 × 1 μs) MD simulation trajectories.

To further dissect these effects, we enumerated persistent interhelical hydrogen bonds and van der Waals (vdW) contacts within the enNTS1 transmembrane region (>40% contact frequency; Figure 2c-d). In apo simulations, hydrogen bonds increased modestly from DM (321) to DDM (325) to LMNG (330). Upon NT(8-13) binding, bond counts rose to 340 in DM and 354 in both DDM and LMNG. For vdW contacts, apo enNTS1 exhibited 939 (DM), 938 (DDM), and 969 (LMNG). With ligand bound, these increased to 993 (DM), 1002 (DDM), but only 979 (LMNG). Together, these data suggest that ligand-induced stabilization arises primarily from vdW contacts in DM and DDM, but is dominated by hydrogen bond formation in LMNG. Thus, detergent environments not only modulate receptor baseline stability but also reshape the mechanism of ligand-induced stabilization. In LMNG, extensive detergent packing reduces the conformational flexibility available for ligand binding to stabilize, leading to smaller ΔG_ligand_ contributions compared to DM or DDM.

### Micelle shape reflects detergent headgroup chemistry

To examine how detergent chemistry influences the global geometry of enNTS1 proteomicelles, we analyzed micelle eccentricity from the most populated conformation (i.e. the largest cluster from cluster analysis performed for the whole proteomicelle) of our MD simulation (Figure 3a,b)^41,42^. The receptor–detergent complexes adopted spheroidal shapes whose eccentricity varied with detergent composition. In the apo state, LMNG micelles were less oblate than DM or DDM micelles (Figure 3c). This is consistent with micelle packing models showing that larger headgroups disfavor oblate shapes^43,44^. LMNG’s branched headgroup likely introduces electrostatic repulsion that promotes a more spherical geometry. Upon NT(8-13) binding, LMNG proteomicelles shifted toward more oblate conformations, resembling DM and DDM, and displayed a broader population distribution (Figure 3d). These results suggest that ligand binding perturbs micelle geometry most strongly in LMNG, potentially reflecting rearrangements in the transmembrane helical bundle that challenge detergent packing. Thus, micelle shape analysis highlights how detergent headgroup chemistry sets the baseline organization of the proteomicelle, while ligand binding induces compensatory geometric changes.

**Figure 3.**
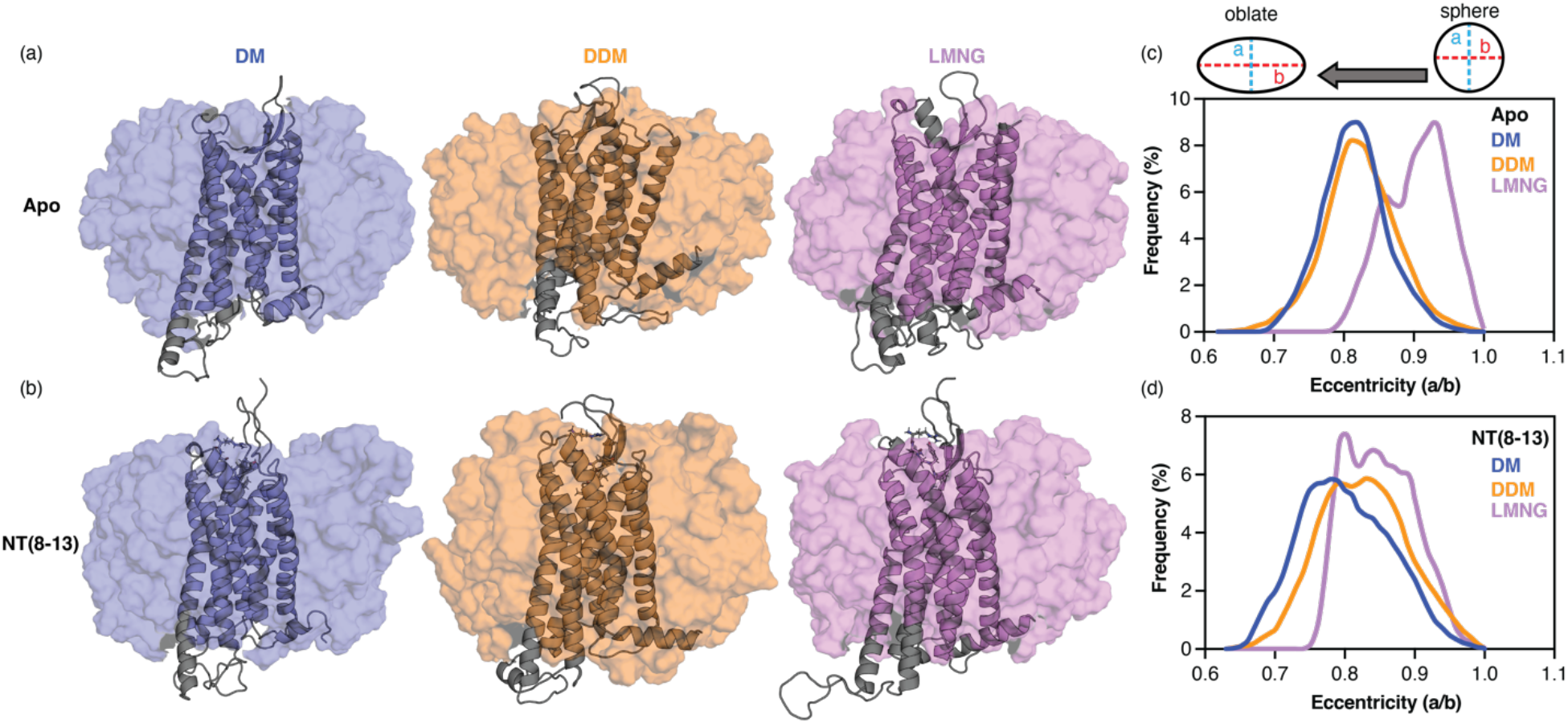
Detergent headgroup chemistry shapes enNTS1 proteomicelles and ligand-induced geometry changes. Representative structures of (a) apo and (b) NT(8-13)-bound enNTS1–detergent proteomicelles from MD simulations. Calculated eccentricity of (c) apo and (d) NT(8-13)-bound enNTS1 in DM (blue), DDM (orange), and LMNG (purple). Eccentricity is defined as the ratio of the short axis over the long axis (a/b). Analysis was performed on the 5 μs (N= 5 × 1 μs) MD simulation trajectories.

### Detergent packing dynamics reveal a trade-off between baseline stability and ligand-induced effects

To evaluate how detergent molecules interact dynamically with the receptor, we examined packing behavior in MD simulations. RMSF measurements showed that LMNG molecules were less flexible than DM or DDM, indicating tighter packing in both apo and NT(8-13)-bound states (Figure S5). Spatial distribution analysis of single detergent molecules further illustrated this effect: DM and DDM diffused more freely, whereas LMNG molecules remained comparatively fixed over time (Figure 4). Radial distribution function (RDF) analysis quantified detergent density within 2 nm of the receptor core (Figure 5a). LMNG exhibited the highest headgroup and tail densities in both apo and holo states, with ∼60% (head) and ∼39% (tail) greater density than DDM in the apo state and ∼51% (head) and ∼39% (tail) increases in the holo state. DM consistently displayed the lowest densities. These results indicate that LMNG forms a more compact and continuous detergent shell around the receptor.

**Figure 4.**
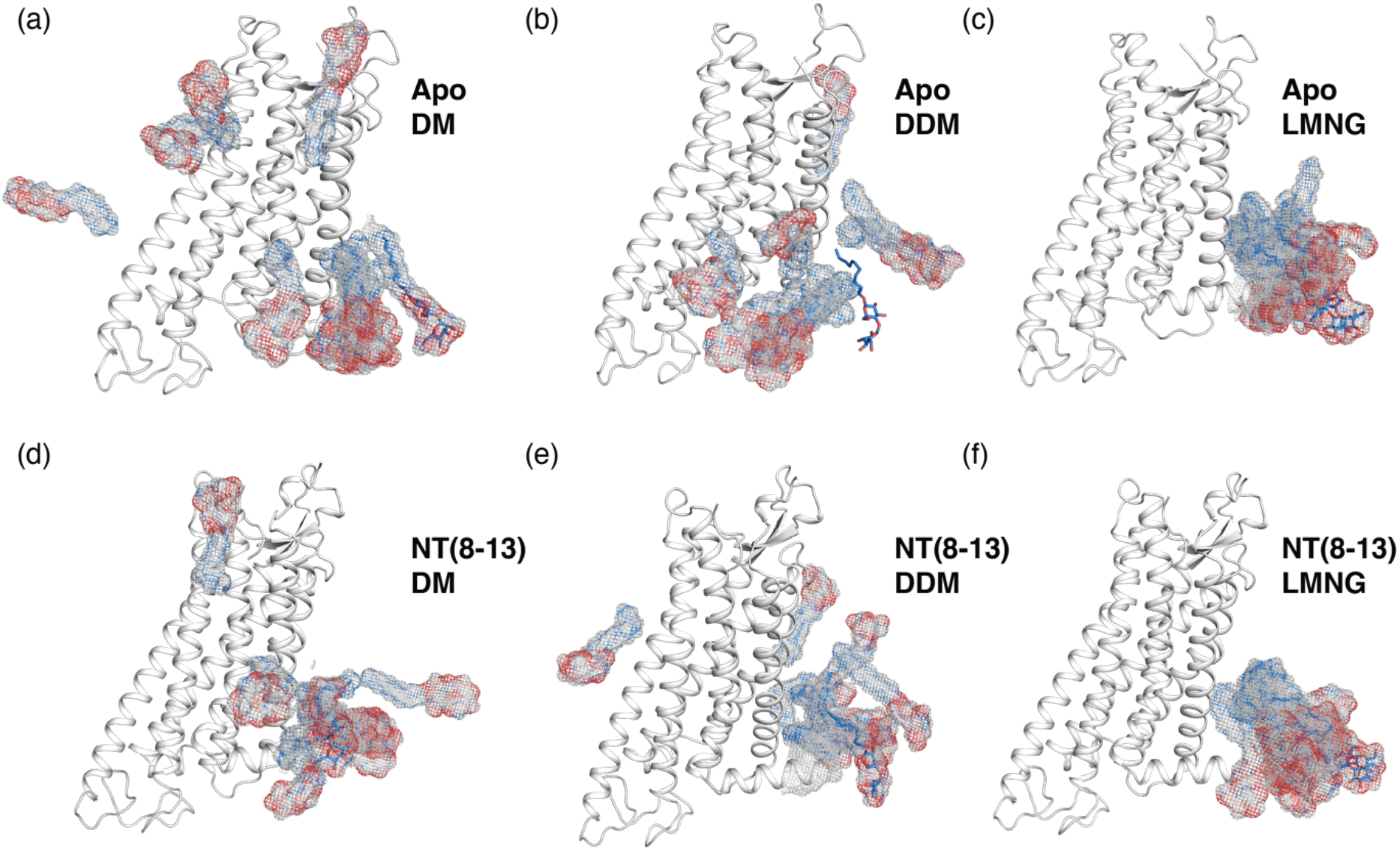
Detergent mobility differs across micelles with LMNG exhibiting reduced dynamics. The spatial distribution function for single (a,d) DM, (b,e) DDM, and (c,f) LMNG molecules are illustrated on representative (a-c) apo and (d-f) holo enNTS1 models. Initial positions of the detergent molecules are shown as stick representations. Spatial distributions are shown as blue (carbon atoms) and red (oxygen atoms) dots. Analysis was performed on the 5 μs (N= 5 × 1 μs) MD simulation trajectories.

**Figure 5.**
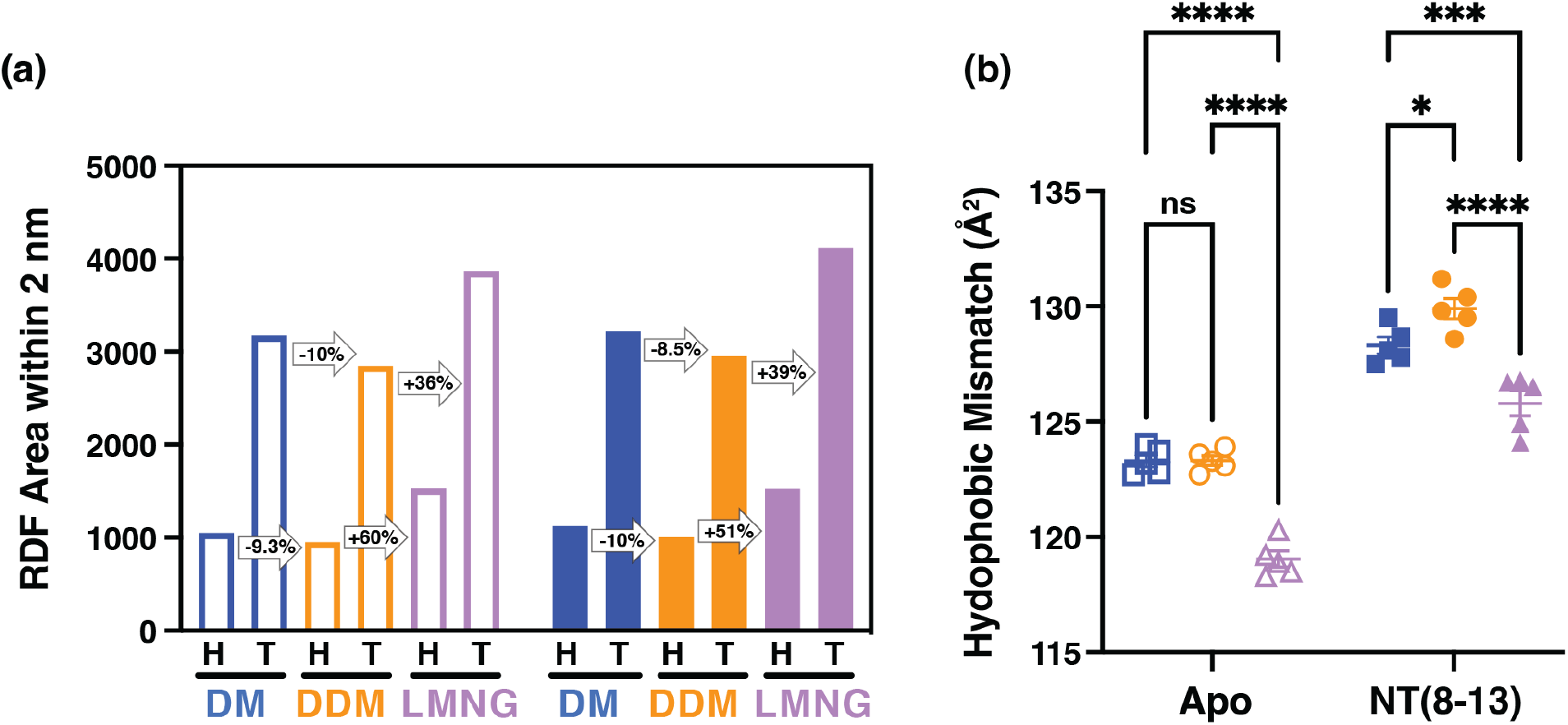
Detergent packing density and hydrophobic mismatch vary between DM, DDM, and LMNG micelles. (a) Plot of the total areas under the radial distribution function (RDF) curves within 2 nm of the receptor core for both the head (H) and tail (T) regions of DM (blue), DDM (orange), and LMNG (purple). The open bars represent the apo state and the filled bars represent the NT(8-13)-bound state. (b) Residual hydrophobic mismatch displaying total surface area of the TM regions exposed to solvent for enNTS1 in DM (blue square), DDM (orange circle), and LMNG (purple triangle) both apo and NT(8-13)-bound. Analysis was performed on the 5 μs (N= 5 × 1 μs) MD simulation trajectories. Analysis was done with two-way ANOVA with multiple comparison using Tukey’s test. (nonsignificant [ns] P>0.05, ^*^P≤0.05, ^***^P≤0.001, ^****^P≤0.0001)

Residual hydrophobic mismatch, which occurs when the hydrophobic thickness of the membrane mimetic does not match the hydrophobic thickness of the membrane protein, supported this interpretation. When taking the non-persistent contacts with less than 20% frequency, DM and DDM exhibited ∼3–4 Å^2^ greater solvent-exposed transmembrane surface area than LMNG, consistent with looser packing (Figure 5b). NT(8-13) binding increased mismatch across all three detergents, but the effect was most pronounced in LMNG, where tighter apo packing was disrupted by ligand-induced conformational changes. Together, these findings demonstrate that LMNG provides superior hydrophobic coverage and the most rigid packing of enNTS1 in the apo state. However, ligand binding destabilizes this optimized packing, leading to smaller relative gains in stability compared to DM or DDM. In thermodynamic terms, LMNG maximizes detergent-driven stabilization (ΔG_conf_) while constraining additional ligand-induced stabilization (ΔG_ligand_).

### Detergent environments redistribute NT(8-13) contacts, with Y11 as a sensitive reporter

MD simulations of agonist-bound enNTS1 in DM, DDM, and LMNG provided insight into how detergent environments reshape ligand dynamics. Analysis of NT(8-13) backbone RMSD revealed overall stability of ∼0.1–0.2 nm across all detergents, but LMNG trajectories showed intermittent fluctuations up to 0.3 nm, including early instability (<200 ns) and later spikes at ∼300, 550, and 700 ns (Figure S7). These results suggest that LMNG alters the conformational ensemble of the bound peptide relative to DM and DDM. Residue-level contact frequency analysis highlighted detergent-dependent differences (Figure S8a). Most strikingly, NT(8-13) residue Y11 exhibited the largest variance in contacts across detergents (Figure S8b), with more modest differences observed for R9 and I12. By contrast, P10 maintained consistent contact frequencies regardless of detergent environment. R9 displayed a slight preference for DM and DDM over LMNG, I12 favored Y347 in LMNG and DM over DDM, and L13 showed enhanced contact with Y351 in LMNG.

The Y11 contact profile was particularly sensitive to detergent chemistry (Figure 6). DM and DDM yielded broadly similar patterns (Figure 6a,d): DM-preferred contacts were located in the N-terminus and ECL1, while DDM-preferred contacts clustered in ECL2. LMNG, however, produced a distinct shift in Y11 interactions. Contacts with L55 (N-terminus), H132 (ECL1), and H348 (helix 7) were reduced, while interactions with S214 in ECL2 became strongly favored (Figure 6b,e,f). Together, these results demonstrate that detergent environments remodel the binding microenvironment of NT(8-13), with Y11 acting as a sensitive reporter of these changes. The similarity of DM and DDM underscores their comparable packing properties, whereas LMNG produces a distinct redistribution of contacts, consistent with its denser detergent shell and reduced conformational heterogeneity. Functionally, this implies that LMNG stabilization (ΔG_conf_) constrains the range of ligand-induced conformational adjustments (ΔG_ligand_), altering how key residues such as Y11 engage the receptor.

**Figure 6.**
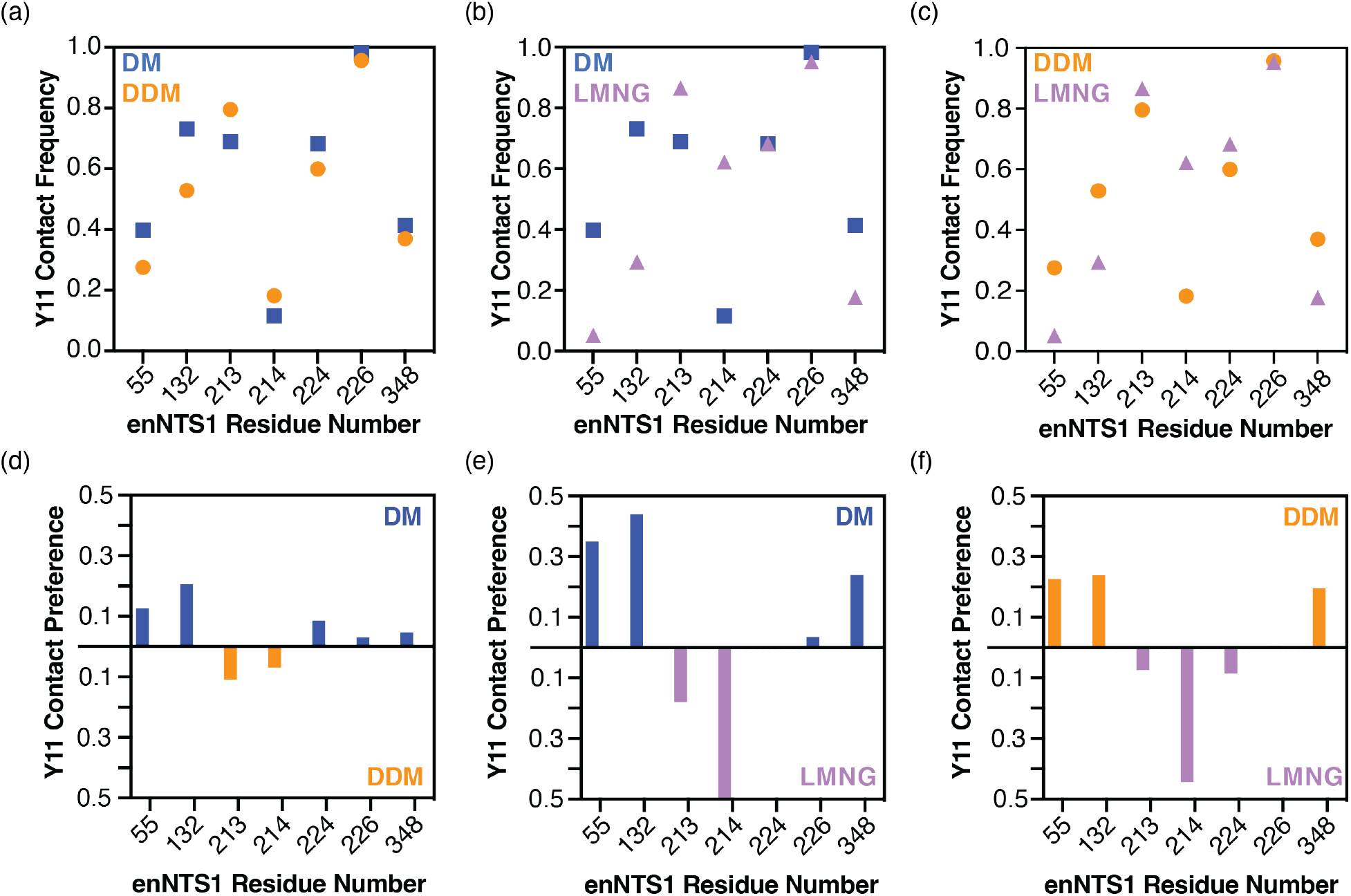
Detergent-dependent redistribution of NT(8-13) Y11 contacts highlights altered binding pocket interactions. Plots of Y11 contact frequency with enNTS1 residues 55, 132, 213, 214, 224, 226, and 348 in (a) DM vs DDM, (b) DM vs LMNG, and (c) DDM vs LMNG. Plotted deltas of Y11 contact frequencies with enNTS1 residues displaying calculated differences between (d) DM and DDM, (e) DM and LMNG, and (f) DDM and LMNG. Analysis was performed on the 5 μs (N= 5 × 1 μs) MD simulation trajectories.

## DISCUSSION

Detergents remain indispensable for GPCR structural biology, but their molecular influence extends beyond solubilization to shaping receptor energetics. Our work shows that detergent chemistry partitions stability into two thermodynamic components: detergent-driven conformational rigidity (ΔG_conf_) and ligand-induced stabilization (ΔG_ligand_). This framework explains the paradox that LMNG, while conferring the greatest baseline stability to enNTS1, simultaneously supports the weakest agonist binding affinity.

In DM, the detergent environment permits broad conformational heterogeneity of the apo receptor, which ligand binding resolves into a more rigid, cooperative state. The large Δ*T*_*m*_ and energetic shifts observed in DM thus reflect substantial ΔG_ligand_ contributions, consistent with the receptor’s engineered background in this detergent^45,46^. DDM provides an intermediate case, where detergent interactions contribute more strongly to stability but still allow significant ligand-induced gains. In contrast, LMNG maximizes ΔG_conf_ as MD simulations revealed dense detergent packing, reduced detergent mobility, and minimized hydrophobic mismatch. As a result, the apo ensemble is rigidified, and ligand binding produces only modest additional stabilization.

This thermodynamic redistribution has functional consequences. Analysis of NT(8-13) dynamics showed that residue Y11, a known reporter of receptor activation^27,28,30^, displayed detergent-specific contact patterns. In DM and DDM, Y11 engaged the N-terminus and ECL1, while in LMNG it preferentially contacted ECL2. Such detergent-dependent remodeling of the binding pocket suggests that LMNG stabilization constrains ligand-induced conformational adjustments, narrowing the energetic landscape available for activation.

Our findings echo broader observations that detergents and membrane mimetics can bias GPCR ensembles ^26,31-33^. While LMNG excels at stabilizing receptors for crystallography and cryo-EM, it may underrepresent conformational states important for signaling or pharmacology. Conversely, less rigid detergents may better preserve ligand-induced transitions but at the cost of reduced overall stability. This balance highlights the need for careful detergent selection depending on whether the experimental goal is to maximize structural resolution or to capture conformational dynamics.

By dissecting detergent-versus ligand-driven stabilization, our study reframes detergent effects as an energetic trade-off rather than a uniform stabilizing influence. This conceptual framework should be broadly applicable to GPCRs and other membrane proteins, providing a basis for rational selection of membrane mimetics and interpretation of stability measurements. Ultimately, understanding how ΔG_conf_ and ΔG_ligand_ are partitioned will be critical for aligning biophysical characterization with the functional roles of membrane proteins in their native environments.

## ACKNOWLEDGEMENTS

The project was funded by Indiana Precision Health Initiative (J.J.Z.), a National Institutes of Health (NIH) T32 fellowship (J.B.B.), and NIH grants R00GM115814 (J.J.Z.) and R35GM143054 (J.J.Z.).

## METHODS

### Reagents

The construct used for expression of enNTS1 in *Escherichia coli* BL21(DE3) cells was generated from a previously published plasmid. ^38^ Thermo Scientific™ HisPur™ Cobalt Resin used in the IMAC purification step was purchased from Fisher Scientific (#89964). All pre-packed affinity resin columns used in the purification of enNTS1 (IMAC, IEX, SEC) were purchased from GE (#45-000-192, #17-5247-01, #29-0452-69). The “MyOne Streptavidin T1” Dynabeads used for functional thermostability assays were purchased from Thermo Fisher (#65601). The fluorescent NT(8-13)^AF647^ was made in house using the dye AF647 from AAT Bioquest (#1833).

### enNTS1 plasmid construct and protein expression

The previously characterized functional variant enNTS1^47^ was used in an expression vector (pDS170) containing an open reading frame encoding an N-terminal maltose-binding protein signal sequence (MBPss), followed by a 10x-Histidine tag, a maltose-binding protein (MBP), a NNNNNNNNNNG linker and a HRV 3C protease site (LEVLFQGP) which were linked via a BamHI restriction site (resulting in additional residues GS) to residue T42 of the receptor. At the C-terminus, T416 of the receptor was linked via a NheI restriction site (resulting in additional AS residues) to an Avi-tag allowing biotinylation, another HRV 3C protease site, a GGSGGS linker, a monomeric ultra-stable green fluorescent protein (muGFP), ^48^ and an additional 10x-Histidine tag. The enNTS1 plasmid was transformed into BL21(DE3) *Escherichia coli* cells and plated on LB agar supplemented with 100 µg/mL carbenicillin and 1% (w/v) glucose at 37 °C overnight. Liquid LB media starter cultures were supplemented with 100 µg/mL carbenicillin and 1% (w/v) glucose then inoculated with several colonies and incubated overnight at 37 °C and 220 RPM. 6 liters of 2xYT media were each supplemented with 100 µg/mL carbenicillin and 0.2% (w/v) glucose were inoculated with 10 mL/L of overnight LB starter culture, and incubated at 37 °C and 220 RPM to an OD600 ≅0.3. The cultures were then cooled to 16 °C. Once each culture reached an OD600 ≅0.7, they were induced with 0.3 mM IPTG and incubated for ∼ 18h at 16 °C and 220 RPM. The cultures were harvested via centrifugation at 5,000 rcf and cell pellets stored at -80 °C. The final amino acid sequence for the purified construct was: GPGSTSESDTAGPNSDLDVNTDIYSKVLVTAIYLALFVVGTVGNGVTLFTLARKKSLQSLQSRVDYYLGSLALSSLLILLFALPVDVYNFIWVHHPWAFGDAGCKGYYFLREACTYATALNVVSLSVERYLAICHPFKAKTLMSRSRTKKFISAIWLASALLSLPMLFTMGLQNLSGDGTHPGGLVCTPIVDTATLRVVIQLNTFMSFLFPMLVASILNTVIARRLTVMVHQAAEQARVSTVGTHNGLEHSTFNMTIEPGRVQALRRGVLVLRAVVIAFVVCWLPYHVRRLMFVYISDEQWTTALFDFYHYFYMLSNALVYVSAAINPILYNLVSANFRQVFLSTLASLSPGWRHRRKKRPTFSRKPNSMSSNHAFSTASGLNDIFEAQKIEWHEGSGL EVLFQ

### enNTS1 protein purification

Cell pellets were solubilized in a *solubilization buffer* (100 mM HEPES, 500 mM NaCl, 20% (v/v) glycerol, 10 mM MgCl2, 10 mM imidazole, pH 8.0) supplemented with 100 mg lysozyme, 1 unit DNAse, 0.2 mM PMSF, and one Roche cOmplete EDTA-free protease inhibitor cocktail tablet. The solution was stirred on ice for 30 minutes before being subjected to sonication on ice: 3 minutes processing time (10 s on, 20 s off) at 35% maximum amplitude. The receptor was then solubilized in detergent with the addition of final concentrations of 1% (w/v) DM detergent, 0.12% (w/v) (CHS), and 0.6% (w/v) CHAPS. The solution was incubated at 4 °C for two hours with stirring. The solution was then centrifuged at 24,424 RCF for 45 minutes. The supernatant containing detergent solubilized enNTS1 was then collected and incubated with TALON resin equilibrated with *equilibration buffer* (25 mM HEPES, 10% (v/v) glycerol, 400 mM NaCl, 0.15% (w/v) DM, pH 8.00) at 4 °C for 45 minutes. Following TALON incubation, the receptor solution was placed into a gravity column to allow flow through of unbound protein. The TALON resin was then washed with *TALON wash #1* (25 mM HEPES, 10% (v/v) glycerol, 500 mM NaCl, 0.15% (w/v) DM, 10 mM Imidazole, 4 mM ATP, 10 mM MgCl2, pH 8.0) followed by *TALON wash #2* (25 mM HEPES, 10% (v/v) glycerol, 350 mM NaCl, 0.05% (w/v) DDM, 10 mM Imidazole, pH 8.0). This second wash step also served as a detergent exchange step from DM to DDM. Following detergent exchange in *TALON wash #2*, enNTS1 was eluted with *TALON elution buffer* (25 mM HEPES, 10% (v/v) glycerol, 500 mM NaCl, 0.05% (w/v) DDM, 350 mM Imidazole, pH 8.0) and incubated with 3 mg of HRV 3C precision protease for 16 h or overnight at 4 °C to cleave MBP and muGFP expression tags. The cleaved enNTS1 was concentrated in a 30 kDa MWCO concentrator via centrifugation at 3,800 RCF and then diluted 10-fold in *SP equilibration buffer* (20 mM HEPES, 10% (v/v) glycerol, 0.05% (w/v) DDM, pH 7.4). This dilution was then split into equal thirds with one batch remaining in DDM, one batch exchanging back into DM, and one batch exchanging into LMNG. Each batch was loaded onto an equilibrated 5 mL SP cation-exchange (CEX) column via a GE AKTA Pure system run at 4 mL/min flow rate. The SP CEX column was washed with *SP wash buffer* (20 mM HEPES, 10% (v/v) glycerol, 300 mM NaCl, 0.03% (w/v) DDM, pH 7.4) until the A280 reading stabilized. The DM batch was then subjected to 30 mL of *DM exchange buffer* (20 mM HEPES, 10% (v/v) glycerol, 100 mM NaCl, 1% (w/v) DM, pH 7.4) at 1 mL/min flow rate. The LMNG batch was treated the same way with a *LMNG exchange buffer* (20 mM HEPES, 10% (v/v) glycerol, 100 mM NaCl, 0.1% (w/v) LMNG, pH 7.4). For elution an equilibrated 1 mL Ni2+-NTA column was attached in-tandem after the 5 mL SP CEX column, and the receptor batches were eluted with the appropriate *SP elution buffer* (20 mM HEPES, 10% (v/v) glycerol, 1 M NaCl, [0.03% (w/v) DDM] or [0.15% (w/v) DM] or [0.01% (w/v) LMNG], 15 mM Imidazole, pH 7.4). The enNTS1 elutions were each concentrated in a separate 30 kDa MWCO concentrator via centrifugation at 3,800 RCF and individually injected onto a GE S200 Increase SEC column equilibrated in the appropriate *SEC buffer* (20 mM HEPES, 150 mM NaCl, [0.03% (w/v) DDM] or [0.15% (w/v) DM] or [0.01% (w/v) LMNG], pH 7.4). Following SEC, the desired enNTS1 fractions for each run were pooled, concentrated to 100-300 µM, and flash-frozen via liquid nitrogen and stored at -80 °C.

### Biolayer interferometry (BLI)

BLI measurements were performed using a Gator Prime 8-channel system (GatorBio) in 96-well plate format. All experiments were carried out at ambient temperature. Ni-NTA biosensors were pre-hydrated for 20 minutes in assay buffer (20 mM HEPES, 300 mM NaCl, [0.03% (w/v) DDM], [0.15% (w/v) DM], or [0.00025% (w/v) LMNG]). Binding was measured by dipping ligand-coated probes into a dilution series of enNTS1 concentrations. For DDM, NT(8-13) was loaded at 50 nM onto Ni-NTA sensors, and enNTS1 ranged from 0-50 nM. For DM, NT(8-13) was loaded at 5 nM, and enNTS1 concentrations ranged from 0-200 nM. For LMNG, NT(8-13) was loaded at 5 nM, and enNTS1 concentrations ranged from 0-120 nM.

BLI traces were collected by acquiring a baseline in assay buffer (200 sec), followed by ligand immobilization (3600 sec). A second baseline was acquired post-immobilization (200 sec), followed by the association phase by dipping the sensor into wells containing enNTS1 (1500 sec). Dissociation was monitored by transferring the sensor back into buffer-only wells (1500 sec). Double referencing was used to minimize background noise. Each protein concentration was paired with a buffer-only reference probe, and its signal was subtracted from its corresponding sample. In all cases, a reference well was included where the ligand-coated sensor was dipped into buffer alone during the association step. Raw data were processed by subtracting the reference well and buffer-only reference probe. The binding curves were analyzed using the quadratic equation that accounts for ligand depletion, single-site binding, and non-specific binding in GraphPad Prism (Version 10.4.0).

### Circular dichroism

For CD measurements, purified enNTS1 was diluted to a final concentration of 5 µM in a *CD buffer* (20 mM HEPES, 150 mM NaCl, [0.03% (w/v) DDM] or [0.3% (w/v) DM] or [0.003% (w/v) LMNG]). For agonist bound enNTS1, a final concentration of 50 µM NT(8-13) peptide was added to the CD buffer. Temperature melting curves were collected on a Jasco J-715 Spectropolarimeter using a 1 mm quartz cuvette, monitoring at 222 nm wavelength, a temperature range of 5 °C to 100 °C, a temperature interval of 0.2 °C, and a response time of 1 s. Data was collected for enNTS1 in each detergent (DDM, DM, and LMNG) in both apo and NT(8-13) bound states. All data were collected in triplicate. The data was normalized between the highest temperature and the lowest temperature points and fit to a 6-parameter Boltzmann equation consisting of an low temperature asymptote, high temperature asymptote, low temperature slope, high temperature slope, *T*_*m*_, and *T*_*m*_ slope. For the LMNG holo condition, the high temperature asymptote was constrained to 0.07 based upon fitted values for the other five conditions.

### Functional thermostability assay

Functional thermostability of enNTS1 was tested by measuring the receptor’s ability to bind a fluorescent NT(8-13)^AF647^ ligand over a thermal gradient. The enNTS1 construct was diluted to 25 nM in *reaction buffer* (20 mM HEPES, 150 mM NaCl, [0.03% (w/v) DDM] or [0.3% (w/v) DM] or [0.003% (w/v) LMNG]). Each detergent condition, as apo and NT(8-13) bound consisted of 36 samples which were then subjected to temperatures ranging from 22 °C to 95 °C (2 °C intervals). For the NT(8-13) bound samples, 250 nM of fluorescent NT(8-13)^AF647^ was added to each sample prior to the thermal melt. After the thermal melt, 250 nM fluorescent NT(8-13)^AF647^ was added to both the apo and holo samples. The enNTS1 construct possesses a C-terminal Avi-tag which was used to immobilize the purified receptor on magnetic Dynabeads with a Streptavidin tag. 3.5 µL of 10 mg/mL Dynabead stock pre-equilibrated with *reaction buffer* was added to the samples after the thermal melt and a Promega (#V8351) 96-well magnetic side strip plate was used to capture the Dynabead-enNTS1 complex for at least 5 minutes. Unbound fluorescent peptide was washed away with reaction buffer and the remaining Dynabead-enNTS1-NT(8-13)^AF647^ complex was taken to a plate reader (BioTek Synergy Neo2) to measure final fluorescence intensity. All data were collected in triplicate. The data was normalized between the highest and lowest temperature points and fit to a 6-parameter Boltzmann equation consisting of an upper asymptote, lower asymptote, upper slope, lower slope, *T*_*m*_, and *T*_*m*_ slope. The upper and lower slopes were constrained to 0.

### Molecular dynamics simulations

MD simulations were performed by utilizing the GROMACS package (version 2021/2022) with the Chemistry HARvard Molecular Mechanics (CHARMM) 36 force field for proteins, n-Decyl-β-d-Maltopyranoside (DM), n-Dodecyl-β-D-Maltopyranoside (DDMs) or Lauryl maltose neopentyl glycol (LMNG) detergent molecules, Na+ CL-ions, and using CHARMM Transferable Intermolecular Potential with 3 Points (TIP3P) water as solvent. The enNTS1 models were generated from the ligand-free state (Protein Data Bank (PDB) ID: 6Z66) or the NT(8-13)-bound rat NTS1 (PDB ID: 6YVR) crystal structures. First, the DARPin crystallization chaperone was removed from both initial crystal structures after which Helix 8 was transposed by aligning the template structures with the NT(8-13)-bound NTS1 mutant without Lysozyme (PDB ID: 4BWB) using Maestro (Schrödinger). Next, the sequences of the generated homology models were adapted to mimic the sequence of the experimental enNTS1 setup, and ICL3 was generated using Prime and further optimized with the refine loops module in Maestro. Lastly, missing side chains and hydrogen atoms were added, protein chain termini were capped with neutral acetyl and methyl amide groups, and histidine-protonated states were assigned. We created the simulation box using the CHARMM–Graphical User Interface (CHARMM-GUI) and positioned the receptor in the micelle using the PPM2.0 web server of the OPM database (orientation of proteins in membranes) using the structure inputs of PDB IDs: 6Z66 and 6YVR for alignment of the TM helices of the protein structure and inserted a pre-equilibrated (B)DM, (B)DDM or (B)LMNG micelle. Final system dimensions of the ligand-free and NT(8-13)-bound enNTS1 (B)DM micelle were respectively: 110 Ȧ by 110 Ȧ by 110 Ȧ, including 192 (B)DM detergent molecules, 35,445 water molecules, 150 mM NaCl; and 113 Ȧ by 113 Ȧ by 113 Ȧ, including 192 (B)DM detergent molecules, 39,586 water molecules, 150 mM NaCl. Final system dimensions of the ligand-free and NT(8-13)-bound enNTS1 (B)DDM micelle were: 110 Ȧ by 110 Ȧ by 110 Ȧ, including 192 (B)DDM detergent molecules, 35,206 water molecules, 150 mM NaCl; and 113 Ȧ by 113 Ȧ by 113 Ȧ, including 192 (B)DDM detergent molecules, 39,364 water molecules, 150 mM NaCl. Final system dimensions of the ligand-free and NT(8-13)-bound enNTS1 (B)LMNG micelle were respectively: 122 Ȧ by 122 Ȧ by 122 Ȧ, including 88 (B)LMNG detergent molecules, 49,957 water molecules, 150 mM NaCl; and 125 Ȧ by 125 Ȧ by 125 Ȧ, including 88 (B)LMNG detergent molecules, 54,745 water molecules, 150 mM NaCl. Next, we minimized all six systems, and equilibrated them using a 1 ns long NVT (constant temperature, constant volume) ensemble and consequently with an NPT (constant temperature, constant pressure) ensemble where we gradually reduced the position restraints from 5 to 0 kcal/mol/Ȧ^2^ with each step being 5 ns long. Since Helix 8 was transposed from a different crystal structure, we performed an extra 50 ns 1 kcal/mol/Ȧ^2^ restraint NPT ensemble on the whole protein except for Helix 8, allowing it to adjust to its new orientation. The final part of equilibration involved a 200 ns long unrestrained NPT simulation before running a total of five independent production MD simulations, each assigned with random velocities and a 1000 ns long. Snapshots were captured every 20 ps, and the entire 1000 ns × 5 runs amounting to 5000 ns of simulation time was used for analysis. We visualized all trajectories using PyMOL (Molecular Graphics System, version 2.0, Schrödinger) and Visual Molecular Dynamics (VMD). The trajectories were analyzed using the GROMACS package (version 2021/2022). The convergence of the MD simulations was confirmed by calculating the change in root mean square deviation of the backbone atom coordinates of the residues in the TM region (Apo: TM1 61-91; TM2 100-130; TM3 139-172; TM4 183-206; TM5 231-271; TM6 299-333; TM7 341-370; NT(8-13)-bound: TM1 61-91; TM2 97-130; TM3 139-172; TM4 183-207; TM5 231-271; TM6 301-333; TM7 341-373) and ligand. All data was analyzed using GraphPad Prism 10 (GraphPad Software, San Diego, California USA).

### Contact analysis of NT(8-13) with enNTS1

To establish contacts between NT(8-13) and enNTS1 in DM, DDM, and LMNG, we used *Get_contacts* (https://getcontacts.github.io). Contact frequencies were calculated from the aggregated trajectories and a contact frequency cut-off of 40% was applied (i.e., counting for at least two MD simulations runs). The frequencies were displayed in a contact heatmap using GraphPad Prism 10 (GraphPad Software, San Diego, California USA).

### Interaction energy of TM backbone of enNTS1

Interaction energies were calculated on the TM backbone of enNTS1 using the *gmx energy* function of the GROMACS package. Both Lennard-Jones short-range and Coulomb short-range energies were calculated on all nonbonded interactions between atoms and summed as the total interaction energies.

### Radial distribution function of detergent molecules

For analyses of how DM, DDM and LMNG molecules organize around the enNTS1, we calculated the radial distribution function for the head and tail of the detergent molecules from the center of mass of receptor using *gmx rdf* function of the GROMACS package.

### Spatial distribution function of detergent molecules

For the localization of DM, DDM and LMNG molecules around the enNTS1 during the MD simulations, we captured the position of one detergent molecule at the beginning and end of each MD run (i.e., 0 and 1000 ns) and these were displayed as a mesh representation around enNTS1 using PyMOL. The initial position of the detergent molecules was shown as stick representation.

### Eccentricity of micelle

The eccentricity of the ellipse of the DM, DDM and LMNG detergents around enNTS1 is defined as the ratio of the long axes to the short axes and was calculated using *gmx gyrate* function of the GROMACS package with the option *-moi*.

### Root-mean-square fluctuation of detergent molecules and TM backbone of enNTS1

The RMSF of the TM-backbone atoms of enNTS1 as well as the detergent molecules DM, DDM, LMNG were calculated using the *gmx rmsf* function in GROMACS. To render the extent of flexibility on the enNTS1 structure as a heat map, we converted the RMSF values to thermal B-factor using loadbfacts in PYMOL.

### Root-mean-square deviation versus α-helicity of TM backbone

The average percent α-helicity of TM backbone residues and RMSD correlation of enNTS1 was obtained by first calculating the α-helicity of every residue in the TM region, using helicity.tcl VMD script which uses STRIDE to identify if a residue is in helical conformation, and subsequently *gmx rms* on all TM-backbone residues (see above). The distributions were plotted using python packages numpy, matplotlib, and seaborn.

### Calculation of hydrophobic mismatch surface area

For calculating unfavorable hydrophobic interactions between enNTS1 and DM, DDM or LMNG, we first identified the hydrophobic residues in the TM that do not make persistent contacts (i.e.,< 20% of the total simulation snapshots) with the detergents tail or head groups using *Get_contacts* (https://getcontacts.github.io). The hydrophobic mismatch surface area between receptor and micelle was obtained by taking the sum of the solvent accessible surface area of TM hydrophobic residues (Glycine, Alanine, Proline, Valine, Methionine, Cysteine, Isoleucine, Tryptophan, Phenylalanine and Tyrosine and Leucine) that show less than 20% persistence contact with DM, DDM, or LMNG (20). The solvent accessible surface area per residue was calculated using the *gmx sasa* function of the GROMACS package.

## REFERENCES

1. Rosenbaum, D.M., Rasmussen, S.G. & Kobilka, B.K. The structure and function of G-protein-coupled receptors. Nature 459, 356 (2009).

2. Venkatakrishnan, A.J. et al. Molecular signatures of G-protein-coupled receptors. Nature 494, 185–194 (2013).

3. Zalewska, M., Siara, M. & Sajewicz, W. G protein-coupled receptors: abnormalities in signal transmission, disease states and pharmacotherapy. Acta Pol Pharm 71, 229–43 (2014).

4. Hauser, A.S., Attwood, M.M., Rask-Andersen, M., Schiöth, H.B. & Gloriam, D.E. Trends in GPCR drug discovery: new agents, targets and indications. Nature Reviews Drug Discovery 16, 829–829 (2017).

5. Isberg, V. et al. GPCRdb: an information system for G protein-coupled receptors. Nucleic Acids Research 44, D356–D364 (2016).

6. Yang, D. et al. G protein-coupled receptors: structure- and function-based drug discovery. Signal Transduction and Targeted Therapy 6(2021).

7. Tate, C.G. Practical Considerations of Membrane Protein Instability during Purification and Crystallisation. in Methods in Molecular Biology 187–203 (Humana Press, 2010).

8. Majeed, S., Ahmad, A.B., Sehar, U. & Georgieva, E.R. Lipid Membrane Mimetics in Functional and Structural Studies of Integral Membrane Proteins. Membranes 11, 685 (2021).

9. Ratkeviciute, G., Cooper, B.F. & Knowles, T.J. Methods for the solubilisation of membrane proteins: the micelle-aneous world of membrane protein solubilisation. Biochemical Society Transactions 49, 1763–1777 (2021).

10. Seddon, A.M., Curnow, P. & Booth, P.J. Membrane proteins, lipids and detergents: not just a soap opera. Biochimica et Biophysica Acta (BBA) - Biomembranes 1666, 105–117 (2004).

11. Chae, P.S. et al. Maltose–neopentyl glycol (MNG) amphiphiles for solubilization, stabilization and crystallization of membrane proteins. Nature Methods 7, 1003–1008 (2010).

12. Lee, H.J., Lee, H.S., Youn, T., Byrne, B. & Chae, P.S. Impact of novel detergents on membrane protein studies. Chem 8, 980–1013 (2022).

13. Caffrey, M. & Cherezov, V. Crystallizing membrane proteins using lipidic mesophases. Nat. Protoc. 4, 706 (2009).

14. Rasmussen, S.G. et al. Structure of a nanobody-stabilized active state of the beta(2) adrenoceptor. Nature 469, 175 (2011).

15. Rosenbaum, D.M. et al. Structure and function of an irreversible agonist-beta(2) adrenoceptor complex. Nature 469, 236 (2011).

16. Granier, S. et al. Structure of the delta-opioid receptor bound to naltrindole. Nature 485, 400 (2012).

17. Haga, K. et al. Structure of the human M2 muscarinic acetylcholine receptor bound to an antagonist. Nature 482, 547 (2012).

18. Kruse, A.C. et al. Structure and dynamics of the M3 muscarinic acetylcholine receptor. Nature 482, 552 (2012).

19. Manglik, A. et al. Crystal structure of the mu-opioid receptor bound to a morphinan antagonist. Nature 485, 321 (2012).

20. White, J.F. et al. Structure of the agonist-bound neurotensin receptor. Nature 490, 508 (2012).

21. Kruse, A.C. et al. Activation and allosteric modulation of a muscarinic acetylcholine receptor. Nature 504, 101 (2013).

22. Ring, A.M. et al. Adrenaline-activated structure of beta2-adrenoceptor stabilized by an engineered nanobody. Nature 502, 575 (2013).

23. Miller, P.S. & Aricescu, A.R. Crystal structure of a human GABAA receptor. Nature 512, 270 (2014).

24. Lee, S. et al. How Do Short Chain Nonionic Detergents Destabilize G-Protein-Coupled Receptors? Journal of the American Chemical Society 138, 15425–15433 (2016).

25. Lee, S. et al. How Do Branched Detergents Stabilize GPCRs in Micelles? Biochemistry 59, 2125–2134 (2020).

26. Zhang, X., Stevens, R.C. & Xu, F. The importance of ligands for G protein-coupled receptor stability. Trends Biochem Sci 40, 79–87 (2015).

27. Cong, X., Fiorucci, S. & Golebiowski, J. Activation Dynamics of the Neurotensin G Protein-Coupled Receptor 1. Journal of Chemical Theory and Computation 14, 4467–4473 (2018).

28. Bumbak, F. et al. Conformational Changes in Tyrosine 11 of Neurotensin Are Required to Activate the Neurotensin Receptor 1. ACS Pharmacol Transl Sci 3, 690–705 (2020).

29. Scott, D.J. & Pluckthun, A. Direct molecular evolution of detergent-stable G protein-coupled receptors using polymer encapsulated cells. J Mol Biol 425, 662–77 (2013).

30. Asadollahi, K. et al. Unravelling the mechanism of neurotensin recognition by neurotensin receptor 1. Nature Communications 14, 8155 (2023).

31. Chung, K.Y. et al. Role of Detergents in Conformational Exchange of a G Protein-coupled Receptor. J. Biol. Chem. 287, 36305 (2012).

32. Rasmussen, S.G.F. et al. Crystal structure of the β2 adrenergic receptor–Gs protein complex. Nature 477, 549–555 (2011).

33. Thomas, M.J. et al. Lipid exchange between mixed micelles of phospholipid and triton X-100. Biochimica et Biophysica Acta (BBA) - Biomembranes 1417, 144–156 (1999).

34. Bumbak, F. et al. Ligands selectively tune the local and global motions of neurotensin receptor 1 (NTS(1)). Cell Rep 42, 112015 (2023).

35. Vogt, G. & Argos, P. Protein thermal stability: hydrogen bonds or internal packing? Folding and Design 2, S40–S46 (1997).

36. Dubois, J.-M., Ouanounou, G. & Rouzaire-Dubois, B. The Boltzmann equation in molecular biology. Progress in Biophysics and Molecular Biology 99, 87–93 (2009).

37. Deluigi, M. et al. Complexes of the neurotensin receptor 1 with small-molecule ligands reveal structural determinants of full, partial, and inverse agonism. Science Advances 7, eabe5504 (2021).

38. Egloff, P. et al. Structure of signaling-competent neurotensin receptor 1 obtained by directed evolution in <i>Escherichia coli</i>. Proceedings of the National Academy of Sciences 111, E655–E662 (2014).

39. Cheng, X., Jo, S., Lee, H.S., Klauda, J.B. & Im, W. CHARMM-GUI micelle builder for pure/mixed micelle and protein/micelle complex systems. J. Chem. Inf. Model. 53, 2171 (2013).

40. Jo, S., Kim, T., Iyer, V.G. & Im, W. CHARMM-GUI: a web-based graphical user interface for CHARMM. J. Comput. Chem. 29, 1859 (2008).

41. Lipfert, J., Columbus, L., Chu, V.B., Lesley, S.A. & Doniach, S. Size and shape of detergent micelles determined by small-angle X-ray scattering. J. Phys. Chem. B 111, 12427 (2007).

42. Oliver, R.C. et al. Dependence of micelle size and shape on detergent alkyl chain length and head group. PLoS One 8, e62488 (2013).

43. Iyer, J. & Blankschtein, D. Are ellipsoids feasible micelle shapes? An answer based on a molecular-thermodynamic model of nonionic surfactant micelles. J Phys Chem B 116, 6443–54 (2012).

44. Dupuy, C. et al. Small Angle X-ray and Neutron Scattering Study of the Micellization of (N-Alkylamino)-1-deoxylactitols in Water. Langmuir 12, 3162–3172 (1996).

45. Sarkar, C.A. et al. Directed evolution of a G protein-coupled receptor for expression, stability, and binding selectivity. Proceedings of the National Academy of Sciences 105, 14808–14813 (2008).

46. Shibata, Y. et al. Thermostabilization of the Neurotensin Receptor NTS1. Journal of Molecular Biology 390, 262–277 (2009).

47. Bumbak, F. et al. Optimization and 13CH3 methionine labeling of a signaling competent neurotensin receptor 1 variant for NMR studies. Biochimica et Biophysica Acta (BBA) - Biomembranes 1860, 1372–1383 (2018).

48. Scott, D.J. et al. A Novel Ultra-Stable, Monomeric Green Fluorescent Protein For Direct Volumetric Imaging of Whole Organs Using CLARITY. Scientific Reports 8, 667 (2018).

