## Supplemental Information for "Stabilization versus flexibility: detergent-dependent trade-offs in neurotensin receptor 1 GPCR ensembles"

**Supplementary Material**

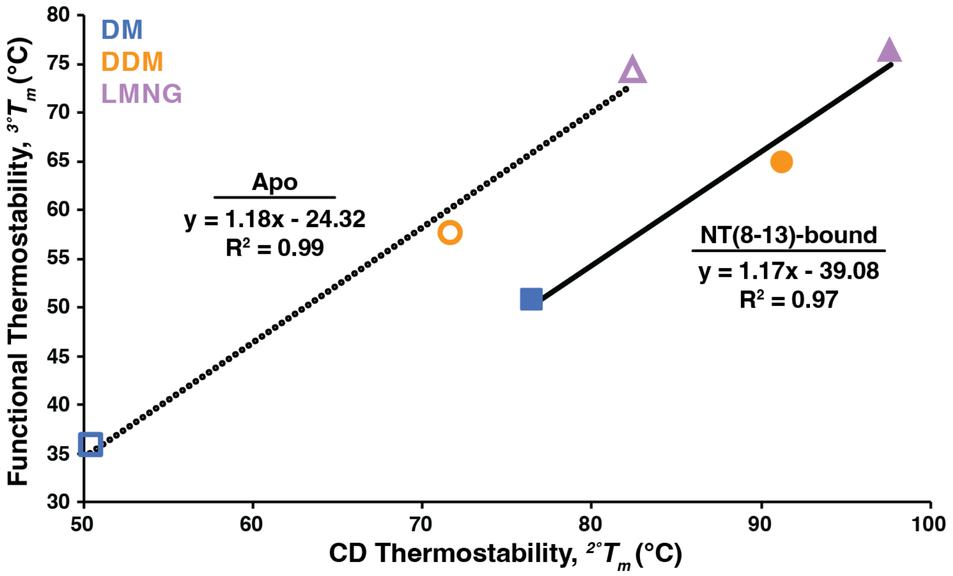


**Figure S1. Correlation between secondary and tertiary thermostability highlights detergent- and ligand-dependent stabilization.** Correlation plot comparing CD thermostability ($^{2^\circ}T_m$) and functional thermostability ($^{3^\circ}T_m$) measurements for enNTS1 in DM (blue square), DDM (orange circle), and LMNG (purple triangle) detergent conditions. Linear fits were applied to both the apo (open shapes, dashed line) and NT(8-13)-bound (filled shapes, solid line) states. Linear fit equation and $R^2$ values are displayed next to each fit. Error bars are smaller than the shapes.

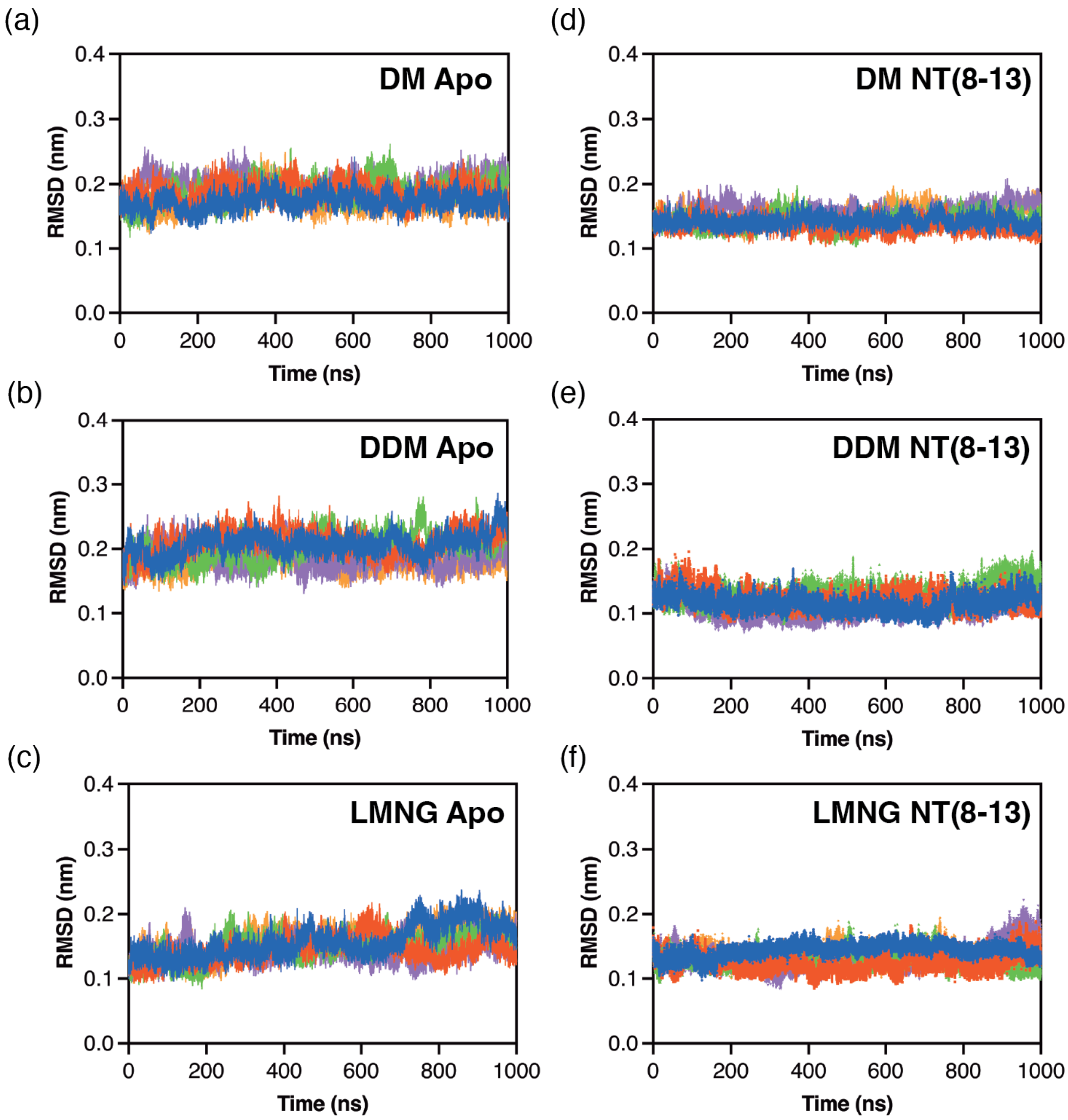


**Figure S2. Detergent environments differentially stabilize enNTS1 transmembrane helices.** TM backbone RMSD values plotted as a function of simulation time for apo enNTS1 model in (a) DM, (b) DDM, and (c) LMNG; and the NT(8-13)-bound enNTS1 model in (d) DM, (e) DDM, and (f) LMNG. Analysis was performed on the 5 μs (N= 5 × 1 μs) MD simulation trajectories.

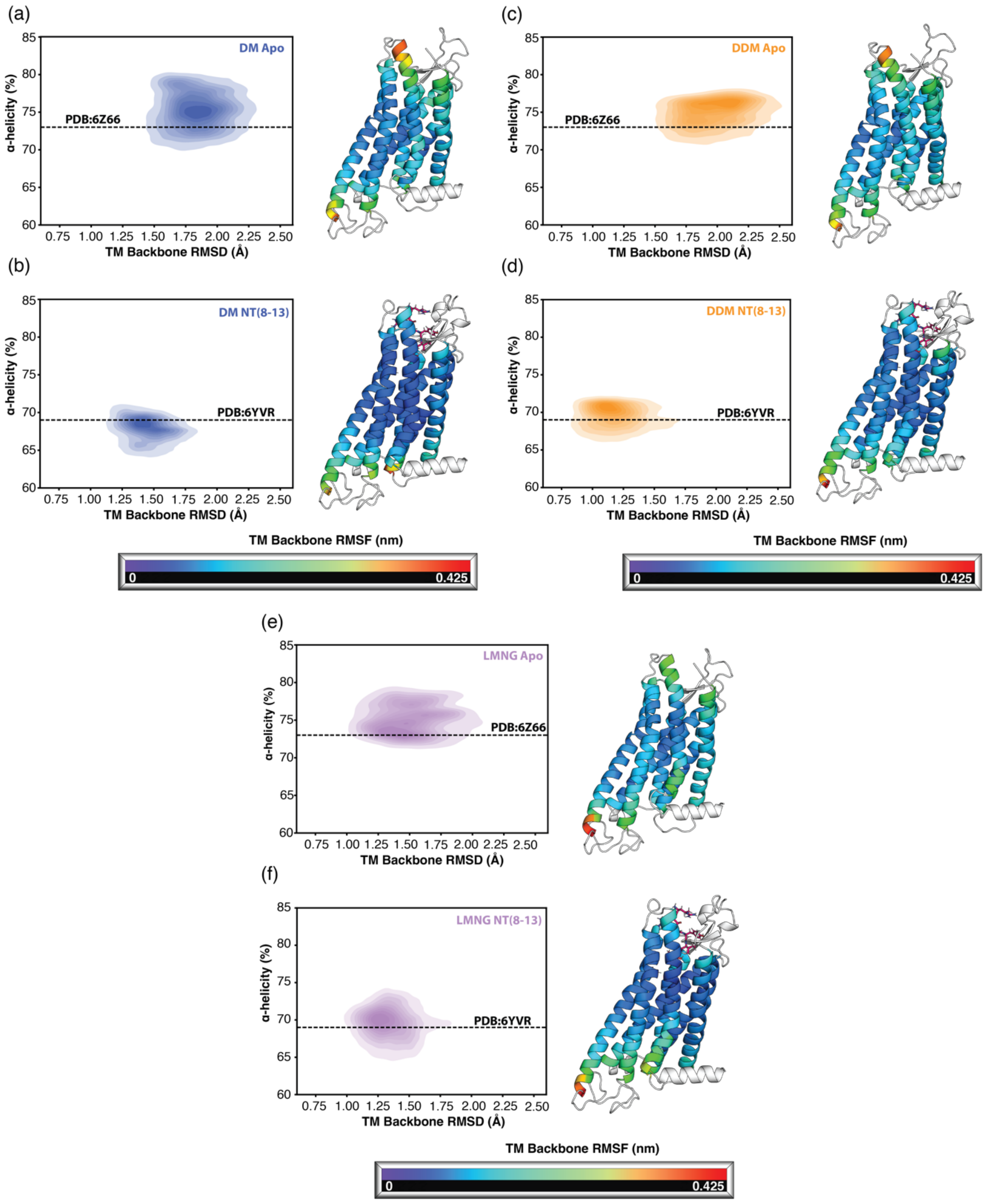


**Figure S3. Conformational heterogeneity of enNTS1 ensembles reveals detergent- and ligand-dependent differences.** Distributions of the average percent helicity of TM region residues and RMSD correlation for each MD snapshot of enNTS1 in DM (a) apo and (b) NT(8-13)-bound, DDM (c) apo and (d) NT(8-13)-bound, and LMNG (e) apo and (f) NT(8-13)-bound. The dashed lines in each plot show the average helicity of the crystal structure used for apo (6Z66) and NT(8-13) (6YVR) enNTS1. Representative structures for each condition are shown with a RMSF heat map of the TM residues. Analysis was performed on the 5 μs (N= 5 × 1 μs) MD simulation trajectories.

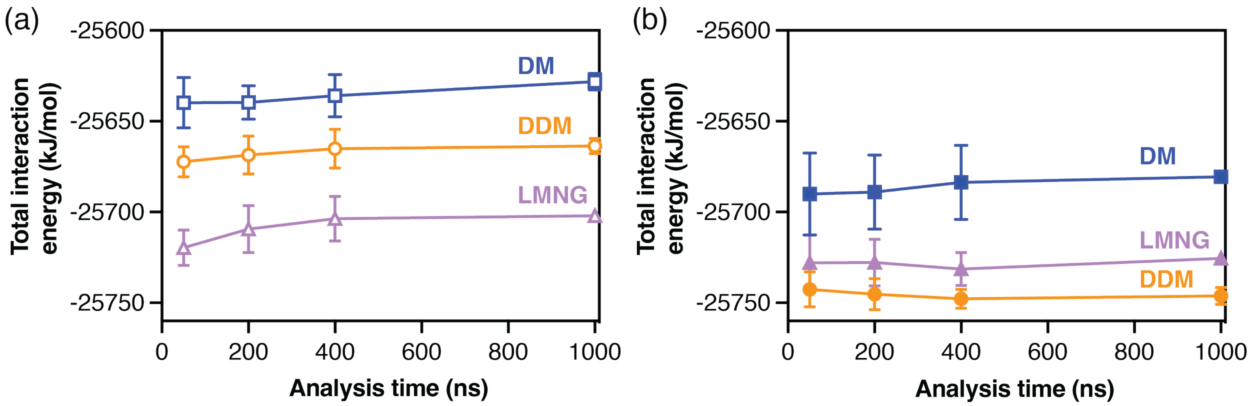


**Figure S4. Detergent-dependent receptor–micelle interaction energies mirror experimental stability trends.** Plots of MD calculated total interaction energies for enNTS1 in DM (blue square), DDM (red circle), and LMNG (green triangle) across the simulation trajectory at 50, 200, 400, and the full 1000 ns. Calculations were done in both the (a) apo and (b) NT(8-13)-bound states. Analysis was performed on the 5 µs (N= 5 × 1 µs) MD simulation trajectories.

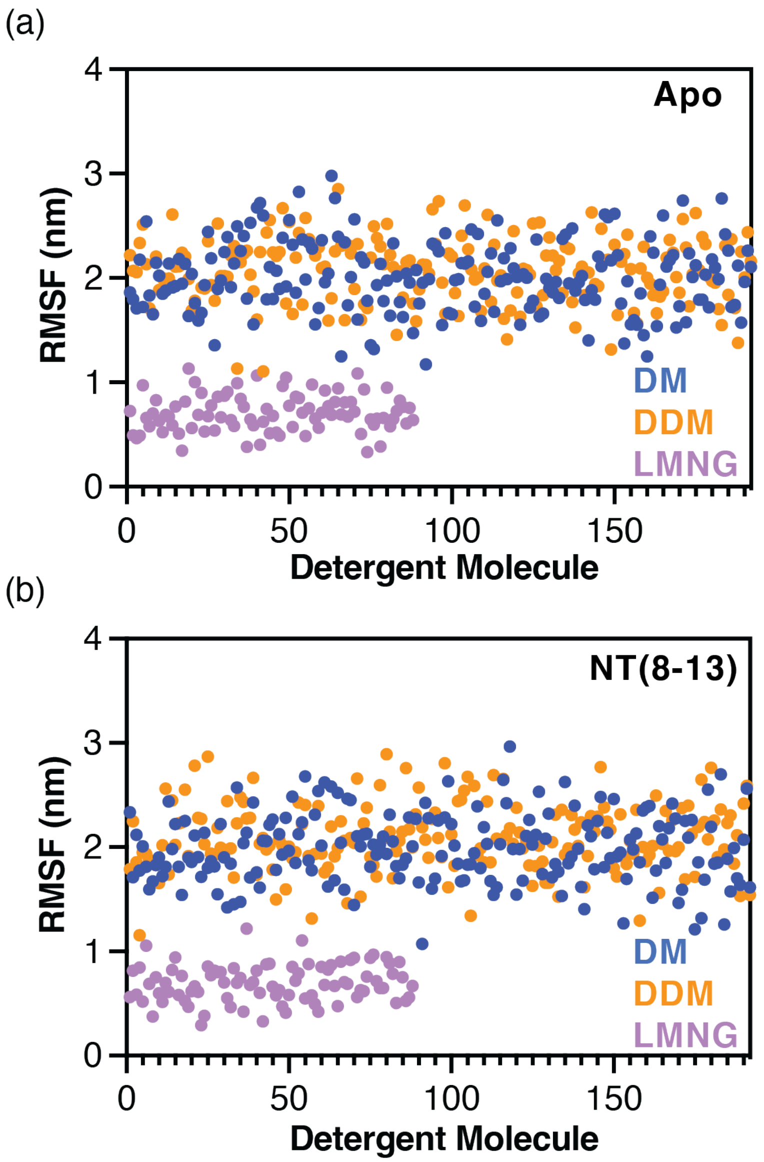


**Figure S5. Detergent mobility differs across micelles, with LMNG forming the most rigid shell.** RMSF of each detergent molecule for DM (blue), DDM (orange), and LMNG (purple) around (a) apo and (b) NT(8-13)-bound enNTS1. Analysis was performed on the 5 μs (N= 5 × 1 μs) MD simulation trajectories.

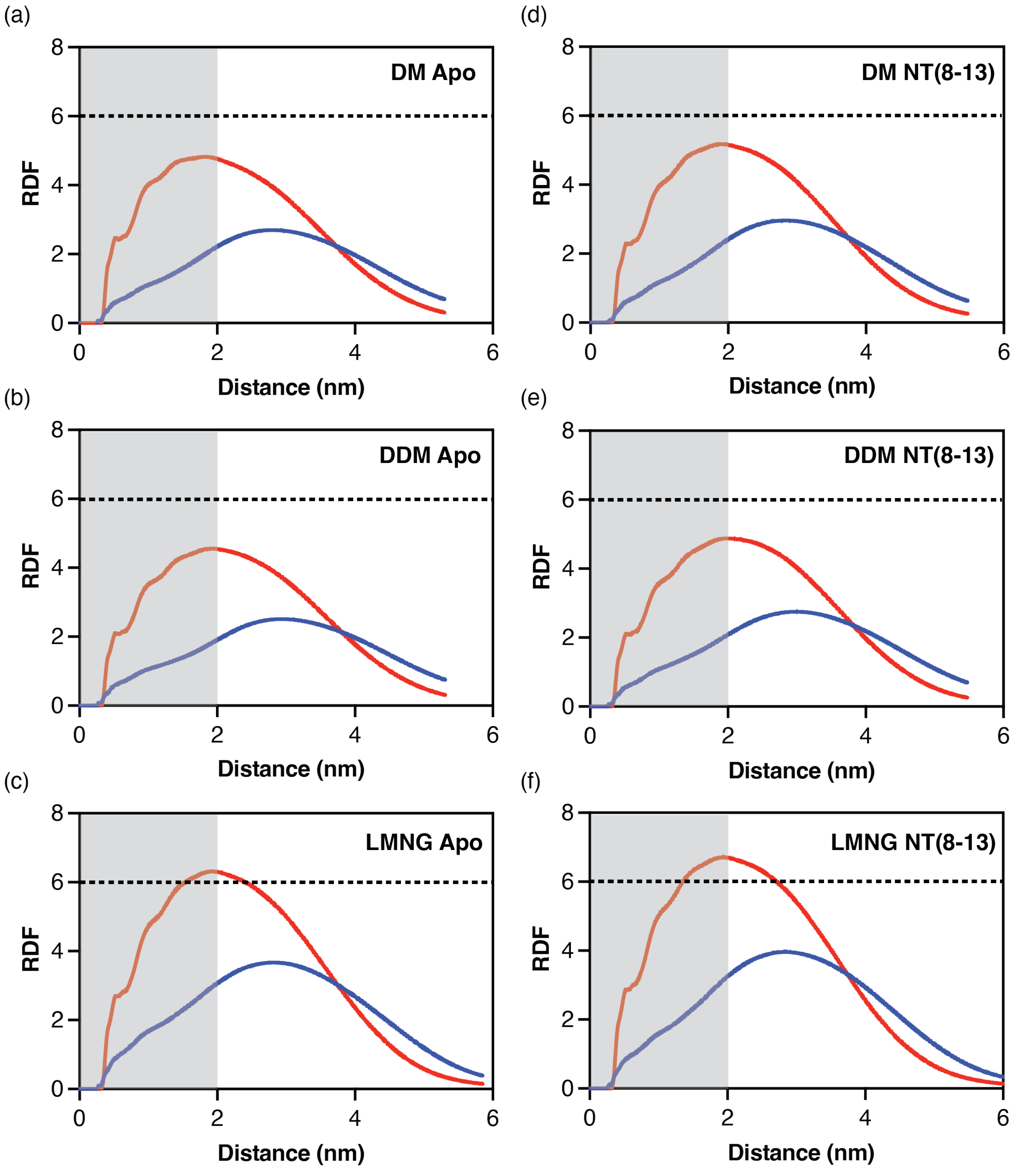


**Figure S6. Radial distribution functions show denser LMNG packing around enNTS1 compared to DM and DDM.** Radial distribution function (RDF) plots displaying the density for either the tail group (red) or head group (blue) of the detergents as a function of distance from apo enNTS1 in (a) DM, (b) DDM, and (c) LMNG; and NT(8-13)-bound enNTS1 in (d) DM, (e) DDM, and (f) LMNG. The gray shaded region represents the first 2 nm from enNTS1. The dashed line is set at an RDF value of 6 for clarity. Analysis was performed on the 5 μs (N= 5 × 1 μs) MD simulation trajectories.

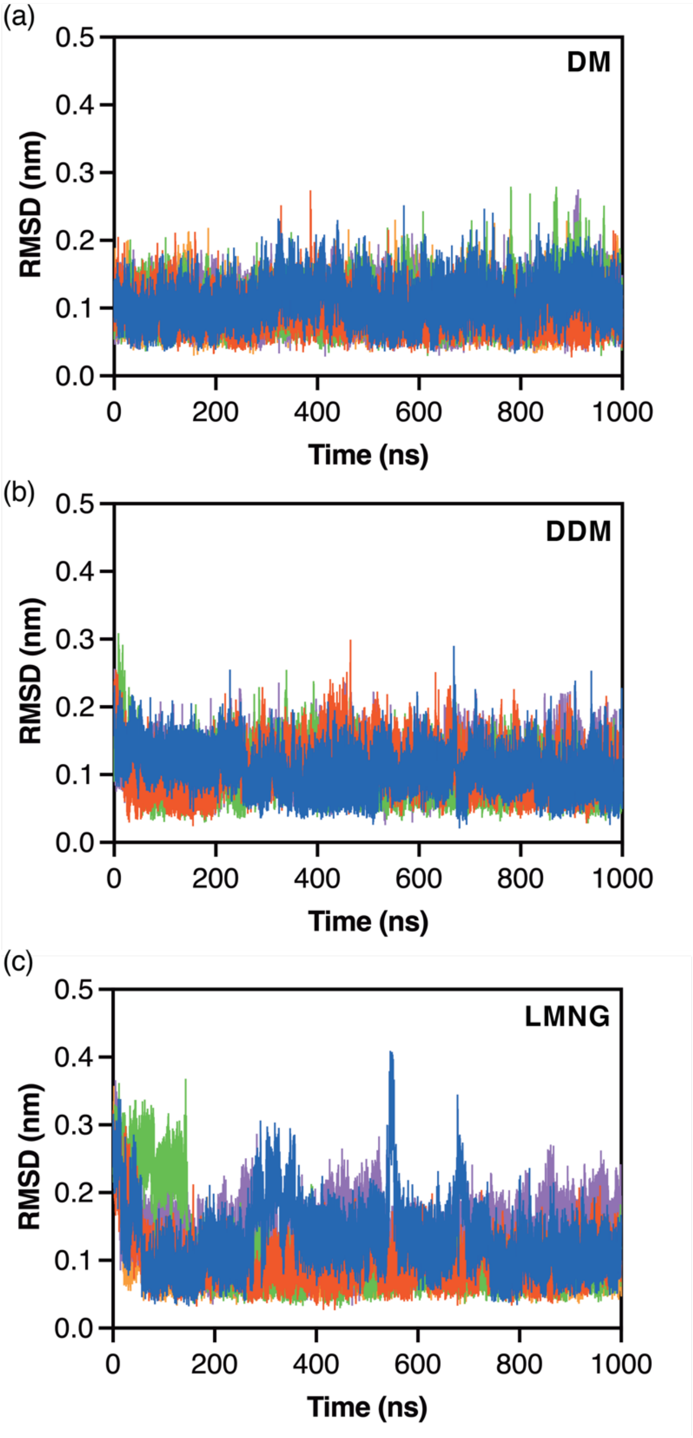


**Figure S7. Ligand backbone dynamics differ by detergent, with LMNG showing intermittent fluctuations.** RMSD values plotted as a function of simulation time for NT(8-13) bound to the enNTS1 model in (a) DM, (b) DDM, and (c) LMNG. Analysis was performed on the 5 µs (N= 5 × 1 µs) MD simulation trajectories.

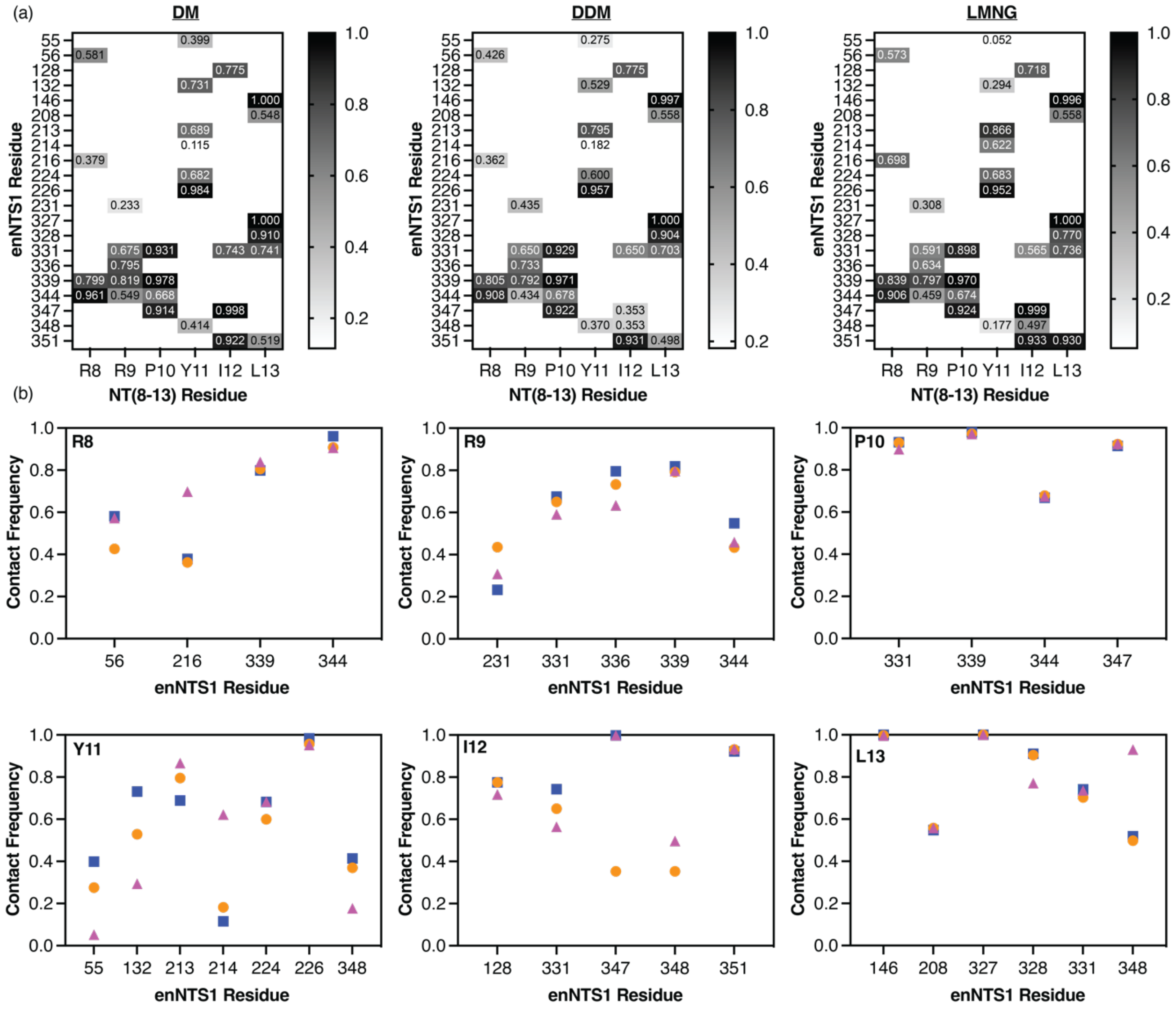

**Figure S8. Detergent environments redistribute NT(8-13) contacts, with Y11 as a sensitive reporter of binding pocket changes.** (a) Contact frequency measurements for NT(8-13) residues with enNTS1 residues in DM, DDM, and LMNG. (b) Comparison of individual NT(8-13) contact frequencies with enNTS1 residues in DM (blue square), DDM (orange circle), and LMNG (purple triangle). Analysis was performed on the 5 μs (N= 5 × 1 μs) MD simulation trajectories.
